# Alevin-fry-atac enables rapid and memory frugal mapping of single-cell ATAC-seq data using virtual colors for accurate genomic pseudoalignment

**DOI:** 10.1101/2024.11.27.625771

**Authors:** Noor Pratap Singh, Jamshed Khan, Rob Patro

## Abstract

Ultrafast mapping of short reads via lightweight mapping techniques such as pseudoalignment has significantly accelerated transcriptomic and metagenomic analyses, often with minimal accuracy loss compared to alignment-based methods. However, applying pseudoalignment to large genomic references, like chromosomes, is challenging due to their size and repetitive sequences.

We introduce a new and modified pseudoalignment scheme that partitions each reference into “virtual colors…. These are essentially overlapping bins of fixed maximal extent on the reference sequences that are treated as distinct “colors” from the perspective of the pseudoalignment algorithm.

We apply this modified pseudoalignment procedure to process and map single-cell ATAC-seq data in our new tool `alevin-fry-atac`. We compare `alevin-fry-atac` to both `Chromap` and `Cell Ranger ATAC`. `Alevin-fry-atac` is highly scalable and, when using 32 threads, is approximately 2.8 times faster than `Chromap` (the second fastest approach) while using approximately one third of the memory and mapping slightly more reads.

The resulting peaks and clusters generated from `alevin-fry-atac` show high concordance with those obtained from both `Chromap` and the `Cell Ranger ATAC` pipeline, demonstrating that virtual color-enhanced pseudoalignment directly to the genome provides a fast, memory-frugal, and accurate alternative to existing approaches for single-cell ATAC-seq processing. The development of `alevin-fry-atac` brings single-cell ATAC-seq processing into a unified ecosystem with single-cell RNA-seq processing (via `alevin-fry`) to work toward providing a truly open alternative to many of the varied capabilities of `CellRanger`. Furthermore, our modified pseudoalignment approach should be easily applicable and extendable to other genome-centric mapping-based tasks and modalities such as standard DNA-seq, DNase-seq, Chip-seq and Hi-C.

## 1 Introduction

ATAC-seq [7] is a widely-used assay that enables the profiling of open-chomatin regions within the genome. It has a wide variety of applications, such as providing insight into gene regulation through the identification of promoters and enhancers [35], studying cell differentiation and development [28, 20], identification of transcription factors [31] to name a few. As with RNA-seq, it is also now possible to profile chromatin at the level of single cells [8, 9].However, the efficient mapping and processing of single-cell ATAC-seq is a computationally-challenging task.

Currently, three main pipelines/methods are used for this task. One involves aligning reads to the genome using methods such as `BWA-MEM` [22], or `Bowtie2` [19], followed by demultiplexing into cells and then finally using existing general-purpose programs like `SAMtools` [23] and `Picard` [37] to sort and filter the alignments. This pipeline can be quite slow when working with large datasets and can be sped up to some extent by (e.g. replacing one of the above alignment tools with `minimap2` [21] as is used in workflows such as `MAESTRO`[38]).

The second approach, `Cell Ranger ATAC` by 10x genomics, provides end-to-end analysis, from read mapping through downstream tasks like cell clustering using called peaks and counts. It is substantially faster than the traditional pipelines due to optimized components that minimize redundant computation and file I/O. However, there is still much potential for further improvement. Additionally, `Cell Ranger ATAC` is released under a non-free license, that permits any use or modification with only 10x genomics’ own proprietary and patented technologies. The third, and the fastest method is `Chromap` [40], which uses a minimizer-based [29] index to maps reads by generating candidate anchors from minimizer hits and aligning candidate positions to the reference. It further speed up mapping by caching the frequently occuring reads

Some approaches, like `scATAK` and `snATAK` [4], use lightweight mapping methods such as pseudoalignment to single-cell and single-nucleus ATAC-seq data. However, they still rely on traditional read alignment tools to align reads to the genome to produce the accurate genomic mappings necessary to extract peaks, whose sequences are treated as experiment-specific targets for pseudoalignment [5]. While lightweight mapping approaches have proven very successful in providing fast read mapping in transcriptomics [5, 26] and metagenomics [30, 33, 24, 1, 12] without much loss in accuracy compared to alignment-based approaches, no method has yet employed lightweight mapping directly to the genome in the context of single-cell ATAC-seq.

One may expect that such approaches might also help in improving the read mapping efficiency for genomic mapping as well. However, the large reference sequences frequently consisting of repetitive regions in the genome make it difficult to applying pseudoalignment. A *k-*mer can map to many locations within the same reference which may lead to an incorrect determination of the mapping position, even when the specific reference (e.g. chromosome of origin) is determined correctly. These issues can lead both to reporting spurious mapping positions as well as missing the most likely approximate mapping location of a read and can also lead to exacerbated multimapping effects.

Recently, `KMCP` [32] described an approach that uses a modified COBs [3] index to tackle the metagenomic profiling problem. The COBs data structure is an (approximate) presence/absence index based on the matching of some threshold fraction of k-mers between a query and the reference. To reduce the potential for spurious matching based off of genomically distant k-mers, and to ensure that the evidence supporting the existence and abundance of a taxon isn’t due to many reads assigned to only one narrow range of the underlying genome (akin to the strategy employed in `KrakenUniq` [6]), `KMC` divides the metagenomic references into a fixed number of chunks, which themselves (rather than the genomes) become the set of distinct targets represented in the COBs index. Taking the success of such an approach as motivation, we propose a modified pseudoalignment approach for mapping reads to the genome, which partitions the references into “virtual colors” of fixed maximum extent. To determine the set of references (and their approximate positions) from which a read originates, the intersection across the virtual colors to which the *k-*mers read map is performed instead of taking the intersection over the original colors (i.e. references/chromosomes). This also necessitates careful handling of the boundaries between virtual colors (to ensure reads spanning virtual colors are not lost) as well as ensuring that duplicate mappings induced by virtual color overlap are properly identified with only a single position in the original reference space.

### 1.1 Contributions

- We present `alevin-fry-atac`,a novel tool designed for the efficient processing and mapping of single-cell ATAC-seq data. It extends the memory frgual `piscem` [11] index, incorporating a modified pseudoalignment algorithm, based upon the concept of virtual colors, that is tailored for accurate genomic mapping without sacrificing efficiency.
- To further enhance mapping speed, we optimized and improved the streaming querying procedure of `SSHash` [27] (a key component of the `piscem` index) and additionally developed a caching mechanism to speed up search of *k-*mers not amenable to streaming query optimizations. Both of these enhancements leverage insights about the structure of the compacted de Bruijn graph, which is a core component of our index.
- We have enhanced the processing functionalities of `alevin-fry` by incorporating a new, data-adaptive, parallelized, external-memory sorting procedure, which outputs a cell-barcode corrected, deduplicated, and coordinated-sorted `BED` file. The `BED` file contains all necessary information to be directly used with popular peak callers such as `MACS2` [41, 13] and other downstream tools for analyzing single-cell ATAC-seq data.
- We evaluated `alevin-fry-atac` on both simulated and experimental datasets and compared it with the other methods for mapping single-cell ATAC-seq data. We observed a high concordance between the peaks and clusters generated on the mapped fragment file produced by `Cell Ranger ATAC` and `Chromap` and those generated on the mapped fragment file produced by `alevin-fry-atac`. `Alevin-fry-atac` uses 300% less memory compared to `Chromap` while taking 280% less time at 32 threads for preprocessing and mapping single-cell ATAC-seq data.
- We have integrated `Alevin-fry-atac` into the `alevin-fry` ecosystem and `simpleaf` [15], a framework originally designed to simplify single-cell RNA-seq analysis using `alevin-fry`. This integration creates a unified, open-source framework that simplifies and streamlines multi-modal analysis.

## 2 Material and Methods

### 2.1 Preliminaries

The `piscem` [11] index is a modular index that enables efficient retrieval of a set of references (colors), as well as all of the associated positions and orientations for a given *k-*mer *x*. Like the `pufferfish` [2] index that preceded it, the `piscem` index is constructed over the compacted colored reference de Bruijn graph, permitting the same set of operations. However, the `piscem` index is substantially smaller than that of `pufferfish`. For a given set of reference sequences, `piscem` uses `cuttlefish` [18] to efficiently construct the compacted colored reference de Bruijn graph and the associated tiling of the reference by unitigs. It then uses `SSHash` [27] to create a *k-*mer to unitig mapping 𝒦 → 𝒰, which maps each *k-*mer to a unitig, relative offset, orientation triplet (*u*_*k*_, *p*_*k*_, *o*_*k*_). `piscem` builds a dense inverted index to represent the unitig-to-reference tiling 𝒰 → ℛ, which allows recalling, for each unitig the reference sequences, and relative positions and orientations of the unitig’s occurrences in the original reference. The *k-*mer to unitig 𝒦 → 𝒰 and unitig-to-reference 𝒰 → ℛ can be combined to correctly and efficiently determine the mapping position (and orientation) of a *k-*mer in the reference, which might be essential for genome-centric mapping tasks. The `piscem` index can be constructed on different types of reference sequences(genome, transcriptome, metagenome, etc.) and the associated software can currently support the processing and mapping of single-cell RNA-seq and bulk RNA-seq. With the introduction of `alevin-fry-atac`, single-cell ATAC-seq is now also supported. A complete manuscript describing the `piscem` index, the key details of the software implementation, and its applications to various indexing tasks is currently under preparation.

Pseudoalignment is a lightweight mapping procedure that aims to assess the “compatibility” between a read and a collection or references without specifically finding the coordinates of each base of the read in the reference. The term was originally coined in the context of RNA-seq quantification [5] where the transcripts to which a read maps were found by doing an intersection across the colors (transcripts) mapped by a set of selected *k-*mers of a read. Certain characteristics of this mapping process, however, predate the coining ([39], [42]). Several approaches and algorithms for pseudoalignment have been proposed. They vary from how and which set of *k-*mers of a read is selected for mapping to how the colors corresponding to the selected *k-*mers are aggregated to get the final color set. The color set denotes the references to which a read maps [1, 12].

The pseudoalignment strategy that has been employed in the current manuscript is the “hybrid” method described in `Themisto` [1] or the “threshold-union” method described in `Fulgor` [12]. Let ℛ= {*R*_1_, *R*_2_, …, *R*_*N*_} denote the set of reference sequences on which we want to map a query sequence *Q*. Let *K* (*Q*) denote the set of *k-*mers for which a mapping 𝒦 → 𝒰 exists in the index. Given the threshold parameter *τ*(0, 1], the output of pseudoalignment for a read is a subset *R*_*C*_ ⊂ ℛ and the relative read mapping position within a reference, such that each reference in the output *R*_*C*_ appears as a mapped hit (i.e. *k-*mer match) across at least *τ* * |*K*(*Q*) | *k-*mers. The mapping position of the leftmost *k-*mer hit of *Q* to the reference is assigned as the mapping position of the read to reference.

### 2.2 Mapping and Pseudoalignment using virtual colors

The accuracy of compatibility information generated by pseudoalignment may be similar to alignment-based approaches when the reference sequences are comparably short, which is the case for transcriptomics and metagenomics (and when the sequenced sample is devoid of non-indexed references that are sequence-similar to the indexed references [34]). However, when directly applying pseudoalignment for mapping to a genome, where the reference sequences are large, comprising of chromosomes that often contain repetitive sequences, we observe a marked drop-off in mapping accuracy using various traditional pseudoalignment approaches. This can happen due to multiple matching locations for a *k-*mer within a single reference that can lead to false positive and false negative mapping positions during read mapping, as demonstrated in Figure 1 A.

**Fig 1:**
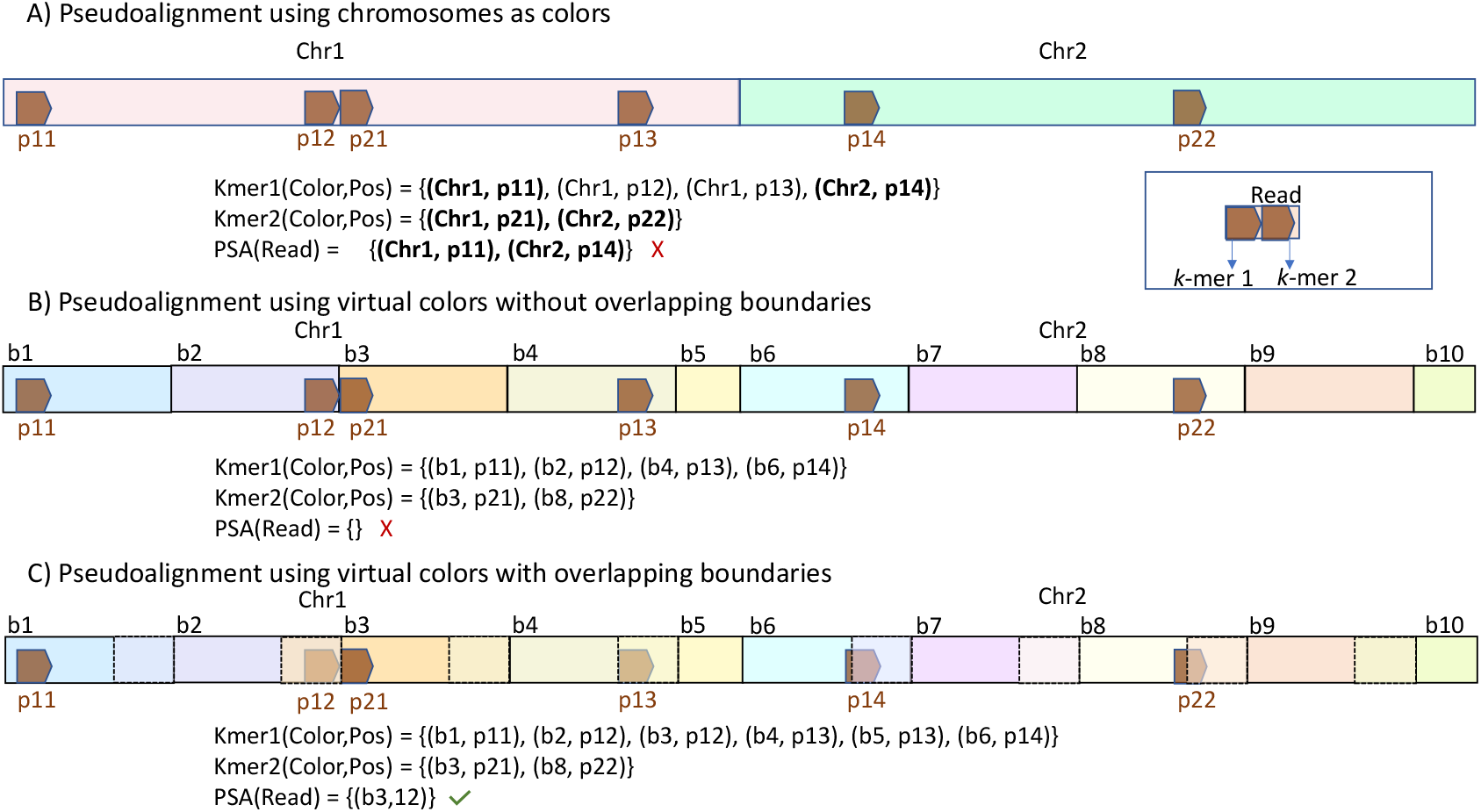
Toy example demonstrating how traditional pseudoalignment on genomic references could lead to spurious mapping or completely missing the correct mapping. In the example, we have two *k-*mers, which map to multiple references, with *k-*mer 1 mapping to multiple locations within chromosome 1. The individual mapped positions within a reference are denoted by p*ij*, with *i* denoting the *k-*mer id and *j* denoting the order of the positions globally for that *k-*mer. PSA(Read) denotes the reference ids and the relative positions on the reference along which Read maps using pseudoalignment. The true mapping for the read in the example is (Chr1, p12). A) Using traditional pseudoalignment both Chr1 and Chr2 are selected as references from which the read originates. The position to which a read maps in pseudoalignment is governed by where the left-most *k-*mer of a read maps. Thus, for reference Chr2, p14 is picked as a position for the hit. If a *k-*mer maps to multiple positions within a reference, then by design the position with the smallest coordinate is picked, which for Chr1 is p11. We thus have two false positive mappings (Chr1, p11), (Chr2, p14) and a false negative since (Chr1, p11) is not reported. B) When the references are projected onto a set of virtual colors whose boundary spaces do not overlap, we miss the mapping if the read spans across the boundaries (b2, b3), creating an Ø set when taking the intersection across the colors. C) When the boundaries of virtual colors overlap, each *k-*mer is assigned two colors if it falls within the overlapping region. This ensures that the read gets mapped after pseudoalignment.

#### Construction of the virtual colors

We propose modifying the pseudoalignment approach by projecting the original reference sequences (i.e. the original colors) onto a set of “virtual colors…, that themselves do not exist as individual elements in the index, but which will represent the targets to which queries can map. The virtual colors, conceptually, consist of a set of overlapping bins, each of fixed maximal extent, on the reference sequences. This overlap is needed since the matching *k-*mers of a read can span the boundary of multiple disjoint virtual colors (and therefore become lost) (Figure 1 B, C). The virtual colors are designed not to extend over the individual reference sequences. Thus a virtual color *b*_*i*_ belonging to reference *R*_*i*_ will not overlap with any virtual color *b*_*j*_ belonging to reference *j*, ⩝*i* ≠ *j*. The construction of the virtual colors is controlled by two parameters namely *𝓁*_vcol_ and *ov_length*. The parameter *ov_length* controls how many bases a virtual color will overlap with the virtual color on its left. It should be at least equal to the read length to ensure that the entire read can fit within it. The parameter *𝓁*_vcol_ controls the total number of bases that will be spanned by a virtual color with most virtual colors on a reference spanning length of (*ov_length* + *𝓁*_vcol_) bases. The leftmost virtual color on a reference will cover a total of *𝓁*_vcol_ bases. The rightmost virtual color on the reference will cover *ov_length* + *rl* bases such that (*B*[*R*_*i*_] 1) *𝓁*_vcol_ + *rl* = *L*[*R*_*i*_], where *B*[*R*_*i*_] denotes the total number of virtual colors contained within the reference sequence *R*_*i*_ and *L*[*R*_*i*_] denotes the length of the reference sequence. *B*[*R*_*i*_] can be computed as 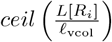. The virtual colors are created dynamically, independent of the index, not creating any overhead.

*Assigning virtual color to a k-mer—* Before starting the mapping process, we create an array *CB* of length | ℛ|, such that:

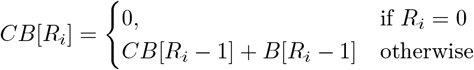

where *CB*[*R*_*i*_] stores the cumulative number of virtual colors up to reference *R*_*i*_. To assign a virtual color to a *k-*mer *x* for which there exists a mapping in the index, we extract the reference id, *R*_*i*_, and 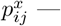 the relative position of the *j*-th occurrence of the *k-*mer *x* within reference *i* in the index. Corresponding to each occurrence of *k-*mer *x* on reference *R*_*i*_. The color id for the *k-*mer *x* is computed as 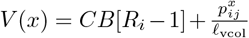. Further, we check if the *k-*mer *x* lies in the overlapping region between the bins. If that is the case, then it is assigned an additional color id *V* (*x*)+ 1. This ensures a read gets mapped if it entirely (or partly) lies within the overlapping regions.

#### Obtaining the final color set

Since a *k-*mer can be assigned two virtual colors corresponding to the same matching position, it might happen that a read after pseudoalignment also gets assigned two virtual colors for that same reference position. This is not a problem for mapping single-end reads, as removing mappings with duplicate color ids and positions is straightforward. However, individual mate mappings must be merged for paired-end reads to get the final set of mapped references and the corresponding starting positions. To do this, we first remove duplicate mappings for the individual mates. Two mappings *a, b* are deemed duplicate : (ori (*a*)= ori (*b*)), (ref (*a*)= ref (*b*)), and (pos (*p*) = pos (*b*)), where ori (·), ref (·), and pos (·) denote the orientation, reference id (original color) and position for a mapped hit, respectively.

By design, a mapped hit will always have *at most one other duplicate* under the above criteria. These hits are deduplicated by keeping the hit with the smaller virtual color id. After removing the duplicate mapping hits for each mate, hits *a* and *b* are merged, with *a* being the mapping for mate 1 and *b* being the mapping for mate 2, if they meet the following criteria :

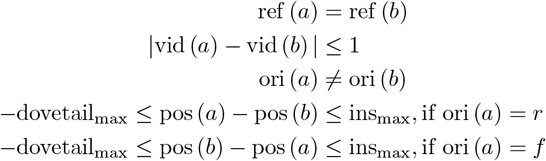

Here vid (·) represents the virtual color id and orientation *f*, *r* represents the forward and reverse orientation respectively. We use the values dovetail_max_ = 20 and ins_max_ = 1000 in this work.

The mapped hits supported by different numbers of *k-*mers can arise when the threshold *τ* < 1. In such cases, we want to pick the hits for which we have the highest confidence, provided by the number of *k-*mer hits. Thus, we keep track of all valid mappings for the read pair, and let *τ*^′^ denote the largest fraction of *k-*mer hits supporting any reported mapping. We subsequently filter out all merged mappings having < *τ*^′^ fraction of valid mapped *k-*mers.

To map the paired-end reads, we first try to merge the individual reads and create a unified fragment. If we are successfully able to merge the reads, then the merged fragment is mapped to the genome instead of the individual reads. We have described this in detail in *Mapping and Pseudoalignment* subsection under Supplementary Methods.

### 2.3 Speeding up k-mer query

A key component of efficiently mapping reads using pseudoalignment is the ability to efficiently retrieve the unitig (and relative offset and orientation within the unitig) to which the *k-*mers in a read map. While the current `piscem` index enables fast extraction of this information, we further optimize the querying procedure to enable even faster retrieval of the unitig and reference information associated with *k-*mers.

#### Streaming queries

Consecutive *k-*mers of read are likely to map to consecutive positions on the reference(s). Consequently, many sequences of adjacent *k-*mers may also originate from the same unitig. Consider, for example, the error (and variation) free case. If the first *k-*mer of a read that is truly drawn from the genome maps to some position *p* in unitig *u*, then the subsequent 𝓁_*u*_ *−p k-*mers will map to the following positions in the unitig. When a *k-*mer is verified to exist in a unitig, we need not search for it anywhere else in the index, since a *k-*mer may appear at most once in the de Bruijn graph. Hence, by optimistically checking for subsequent query *k-*mers that may reside in the same unitig (which, itself, is likely in CPU cache), we can often avoid having to refer to the index entirely until the unitig of origin shifts.

Specificaly, when querying a *k-*mer, we maintain a state variable. This variable stores the information about the unitig to which the previously mapped *k-*mer of the read matched, including its id, orientation, relative offset, and distance from the unitig’s end. The state variable is empty either in two cases: the first *k-*mer of a read is being queried, or the distance of the previous *k-*mer from the end of unitig is 0 (i.e. the last *k-*mer on the unitig w.r.t the orientation of mapping).

In both of these cases, the current *k-*mer is directly queried to the index. Otherwise, the next nucleotide is extracted from the unitig using the previous *k-*mer’s offset and orientation. This nucleotide is matched with the last nucleotide of the current *k-*mer. If they match, then we have the relevant mapping information for this *k-*mer. Otherwise, a query to the index is made. The fewer queries that are made to the main index, the faster the mapping process, the opportunity for which is provided by consecutive *k-*mers in the read. The approach is similar to the streaming iterator in `SSHash` [27]. However, the implementation in `SSHash` tracks the minimizer of the current *k-*mer and queries the corresponding minimizer buckets of the index whenever the minimizer changes, regardless of whether or not a straightforward extension would succeed. In our implementation, by exploiting the structure of the unitigs and the guarantees provided by the de Bruijn graph, we avoid even this lookup, which we find can lead to a substantial (30-40%) improvement in average lookup speed when querying each *k-*mer in a read.

#### Caching terminal k-mers

We aim to exploit the observation that chromatin profile enrichment is confined to specific genomic regions. The mapping information for *k-*mers can be efficiently extracted as described above if the *k-*mers lie within the same unitig. However, a general query in the index is always necessary when querying a *k-*mer that would reside on a different unitig. Whenever the k-mer preceding such a k-mer was the first or last in a unitig, we refer to such *k-*mers as *fk-*mers and implement a caching mechanism to enable efficient retrieval of their mapping information. Specifically, we suspect that mapped reads will repeatedly traverse specific unitig paths in the underlying de Bruijn graph, and so that by inserting the terminal (first or last) *k-*mers in these unitigs in a small and fast cache, we can often avoid lookups in the main index even when moving between unitigs.

The cache is implemented as a shared, concurrent, hash map (we use the `Boost unordered_concurrent_flat_map`) maintained across all mapping threads. Hash map keys are *fk-* mers, and the values are the results of querying the index for those *k-*mers. The map has a fixed size (default 5, 000, 000 entries) and follows a write-once policy, meaning, once a key-value pair is added, it is not replaced.

For each *k-*mer, we first determine if it an *fk-*mer and if so, we check the cache. If the *k-*mer is present in the cache, the corresponding unitig information is retrieved directly. Otherwise, a full query is performed on the index, and if the *k-*mer is found and the map size permits, the *k-*mer is added to the cache.

To identify *fk-*mers, we leverage state information. Specifically, a *k-*mer is labeled as an *fk-*mer if the distance of the previous *k-*mer from the unitig’s end is less than the distance between the offsets of the previous and the current *k-*mers. This policy allows us to avoid even attempting to consult the cache when we expect the next *k-*mer to occur within the same unitig, and to only consult and modify the cache for unitig-terminal *k-*mers (i.e. *fk-*mers).

We note that `Chromap` also employs a cache [40], where the cache is created on a chain of frequently occurring minimizers within a read, storing their corresponding mapping positions from the index. While the idea of using a cache to speed up mapping to a data-dependent subset of the genome is inspired by `Chromap`, the nature of our cache, the elements that are cached, and the manner in which it is implemented, are completely different.

### 2.4 Alevin-fry-atac pipeline

We now describe the `alevin-fry-atac` pipeline for preprocessing single-cell ATAC-seq data. The first step is to map all the reads using the modified pseudoalignment algorithm described above. The default output of this phase is a RAD (reduced alignment data) format file. The RAD format is a chunk-based, binary file optimized for machine parsing that encodes the relevant information necessary for subsequent data processing. Further details of this format are described in [16], though it worth noting that the single-cell ATAC-seq variant of such files carries some distinct information from the single-cell RNA-seq variant (e.g. it includes positions for each mapping, but doesn’t include UMI information).

#### Cell barcode correction

After mapping, the next step is cell barcode correction. While various approaches to barcode detection and correction are possible (`alevin-fry` implements several distinct modes), for `alevin-fry-atac` we currently only support correcting to an unfiltered permit list. The permit list file contains a list of experiment-independent barcodes that are a superset of the barcodes that should be observed in any given sample. We first scan through the RAD file and count the number of times a barcode corresponding to the mapped record appears (exactly). If a cell or barcode has at least `min-reads` records (default 10) mapping to it, and is also in the whitelist, it is marked as present. For all the other mapped records corresponding to cells not currently marked as present, a nearest neighbor search is performed on the barcode against the list of present cells. If a barcode has a unique (i.e. only one) present neighbor at edit distance 1, that barcode is corrected to that of the neighbor. All other records are ignored.

#### Sorting and Deduplication

In this step, a coordinate-sorted BED file is produced. First, a collection of temporary files are created, each corresponding to contiguous ranges of the underlying genome. The input RAD file is parsed and all records are routed to their appropriate temporary file based on their starting mapping location. The temporary files are then individually sorted (in parallel) and the sorted files are merged (in genomic coordinate order) to produce a BED file. During the sorting process, the duplicate entries for a record (i.e., entries with the same reference, start position, end position, and barcode) are collapsed, and the corresponding number of duplicate entries is stored. For each mapped fragment, the BED file contains the reference name, start position, end position, cell barcode and the total number of duplicates (including the fragment itself). The BED file can then be passed to a peak caller such as `MACS2` [13], the output of which, along with the original BED file containing the counts, forms the input for downstream analysis.

### 2.5 Dataset and the analysis pipeline

We evaluated and compared the performance of the different methods on three simulated and three experimental datasets. `Mason` [17] was used to generate the simulations for read lengths 50, 100, 150 bases. One million paired-end fragments for each read length were created with a probability mismatch 0.25, according to the Illumina model in the simulator. The experimental datasets used in this study include the `Human 10K PBMC`, `Mouse 8K Cortex` and `Human 3K Brain` respectively, some of which have been utilized for analysis and benchmarking in previous studies. The datasets have been downloaded from 10X Genomics website, for which the URLs are provided in Table S1. The details regarding the datasets, analysis pipeline, experimental setup, evaluation, benchmarking, and the software that have been used are provided in detail in *Analysis pipeline* subsection under Supplementary Methods.

### 2.6 Simplified execution of alevin-fry-atac using simpleaf

We have also added support for `alevin-fry-atac` to `simpleaf` [15]. `Simpleaf` is an “augmented execution context” originally designed to streamline execution of `alevin-fry` processing of single-cell and single-nucleus RNA-seq data, as well as to improve reproducibility, encode best practices, and simplify metadata propagation. We have added an `atac` subcommand to `simpleaf` which allows running the entire `alevin-fry-atac` pipeline (from mapping through the optional calling of peaks using `macs3` [13]) via a single command. This presents a vastly simplified interface for running `alevin-fry-atac`, and also generates and organizes relevant metadata, encoded in the form of `JSON` files recording key inputs, outputs and parameter choices, into a standard output structure. While the algorithms and methods we have descirbed in this paper represent advancements and new capabilities within both the `piscem` and `alevin-fry` [16] tools, we recommend using `simpleaf` [15] as the standard interface for running `alevin-fry-atac`.

## 3 Results

### 3.1 Mapping accuracy on the simulated data

#### Evaluating the effect of different parameters for alevin-fry-atac

As described in the methods section, `alevin-fry-atac` includes several parameters: the *k-*mer size *k*, pseudoalignment threshold *τ*and *𝓁*_vcol_, which controls the number of bases a virtual color will span (with most virtual colors spanning *𝓁*_vcol_ + *ov_length* bases). The mapping accuracy is closely tied to the value of these parameters. To evaluate this, we generated simulated datasets with varying read lengths (50, 100, 150 base pairs) and on each dataset, tested the accuracy under different combinations of these parameters. For each evaluation, two parameters were held constant while the third was varied. The following parameter values were tested:

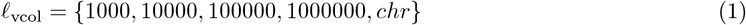

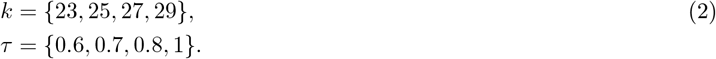

*chr* implies mapping is done using chromosome sequence as colors.

The mapping accuracy for each parameter combination is detailed in Supplementary Results and visualized in Figures S1 - S12. In summary, we observe that virtual colors, regardless of the *𝓁*_vcol_ value, *substantially* enhance the mapping accuracy compared to using chromosomes as colors for pseudoalignment. It has the largest impact on accuracy of all the parameters, where the impact is consistent across different read lengths and parameter combinations. Specifically, the accuracy drops from 93.7% to 84% when the *𝓁*_vcol_ value is changed from 1000 to *chr*, with *k* = 25, *τ*= 0.7 for reads of length 50 (Figure 2).

**Fig 2:**
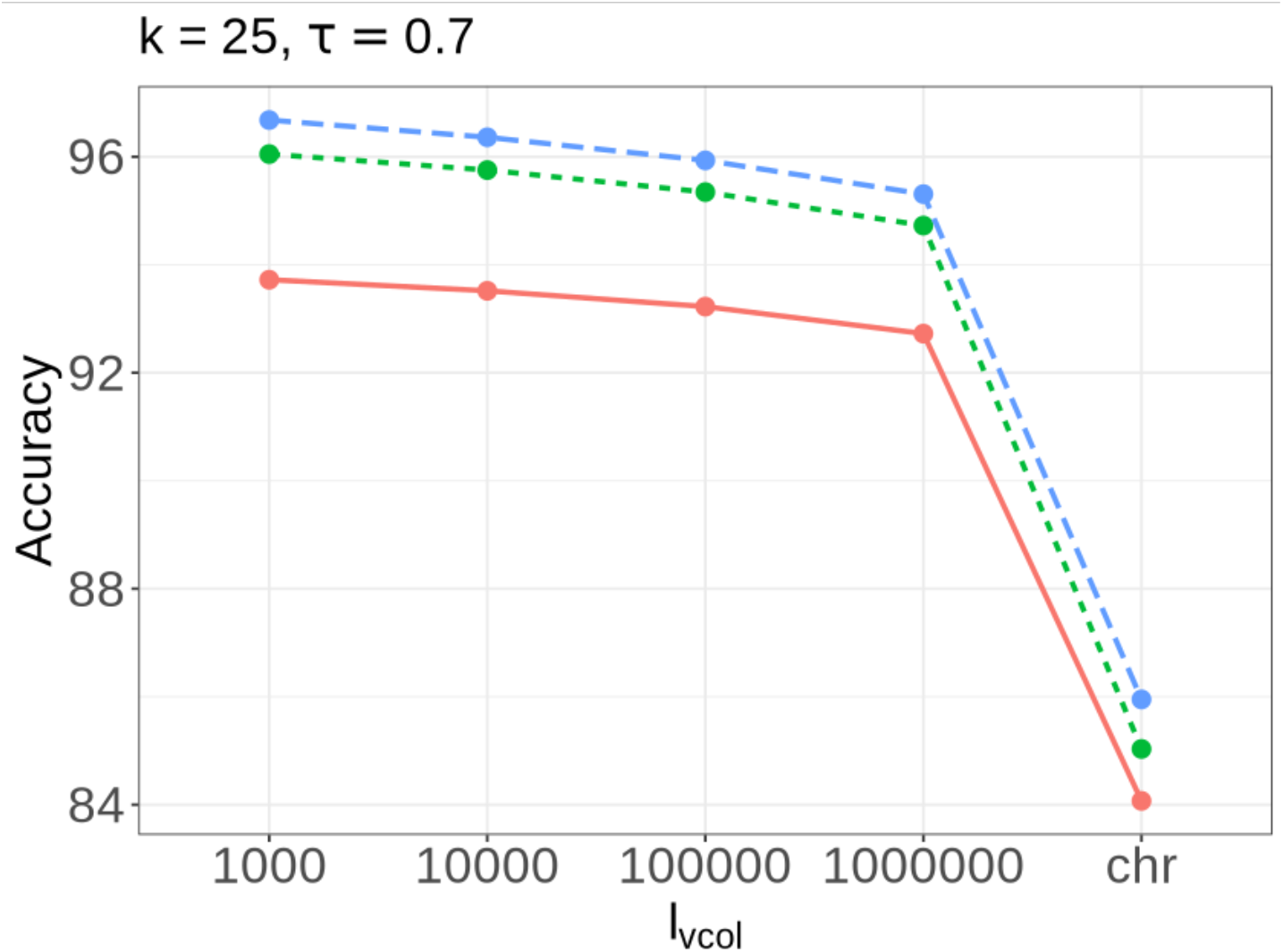
Evaluating accuracy across different read lengths for `alevin-fry-atac` by varying A) *𝓁*_vcol_ for *k* = 25, τ = 0.7. The x-axis label *chr* implies mapping is done using chromosome sequence as colors (removing the impact of virtual colors on mapping procedure).

For reads of length 50, the highest accuracy is observed at *k* = 25, while for reads of length 100 and 150, accuracy improves at larger *k* values. Additionally, accuracy generally decreases as *τ*increases from 0.6 to 1.

#### Comparing the mapping accuracy with the other methods

We next compare the mapping accuracy of `alevin-fry-atac` with `Bowtie2` [19] (which does full read alignment) and `Chromap` on the simulated dataset which is ummarized in Table 1. Similar to `alevin-fry-atac`, the other methods also demonstrate higher accuracy for longer read lengths. Among the methods, `Bowtie2` has the highest accuracy, followed closely by `Chromap`, while `alevin-fry` shows slightly lower accuracy across all the evaluated read lengths. Specifically, for reads of length 50, the mapping accuracy of `alevin-fry` is ∼ 2% lower than `Bowtie2` and around 1% lower for reads of length 150.

**Table 1:**
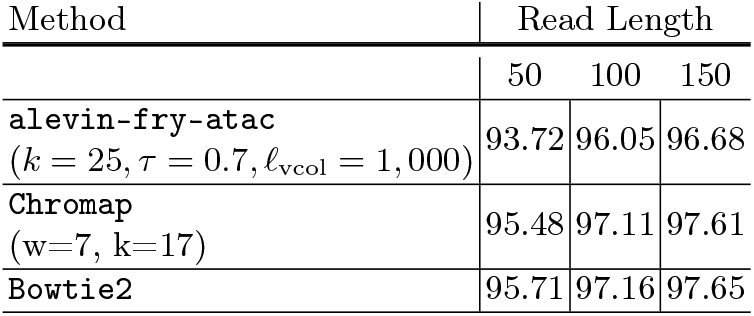
Mapping accuracy on the simulated data for the different methods

### 3.2 Experimental datasets

#### Mapping Rate

We also looked at the impact of varying the different parameters of `alevin-fry-atac` on the overall mapping rate for the different experimental datasets, namely `Human 10K PBMC`, `Mouse 8K Cortex` and `Human 3K Brain` (Figure S13). As with the simulated dataset, we also find a decrease in mapping rate as *𝓁*_vcol_ increases for different *k* and *τ*values, with the fall being the sharpest when chromosomes are used as colors (i.e. when virtual colors are disabled). Further, since we have relatively shorter read lengths in these datasets, higher mapping rate is observed at lower values of *k*. The highest mapping rate is observed for *k* = 25, followed closely by *k* = 23 for the human datasets, while for the mouse dataset the highest mapping rate is achieved at *k* = 23 followed by *k* = 25, with the lowest values observed at *k* = 31 across all datasets. We observe an overall higher mapping rate for `alevin-fry-atac` using *k* = 25, *τ*= 0.7, *𝓁*_vcol_ = 1, 000 (98.27%, 97.73% and 96.99) for the `Human 10K PBMC`, `Mouse 8K Cortex` and `Human 3K Brain` datasets respectively while for `Chromap` (without using the `--preset atac` argument) is (95.31%, 95.56% and 94.50%).

#### Comparing peaks

After obtaining the mapped fragment file—which has been barcode-corrected, deduplicated, and coordinate-sorted—a key step in single-cell ATAC-seq analysis is peak identification. Peaks represent genomic regions with significant enrichment of sequencing reads. To identify peaks, we used `MACS2` on the mapped fragment files generated by `alevin-fry-atac`, `Chromap` and `Cell Ranger ATAC` across different datasets. The total number of bases covered by the peaks is provided in (Table S2), which also includes the bases covered by peaks identified using the custom `Cell Ranger ATAC` peak caller, which could only be run on the `Cell Ranger ATAC` fragment file. Additionally, we did not process the `Human 3K Brain` dataset with `Cell Ranger ATAC`, as it is a multi-omic dataset required analysis with a different software, namely `Cell Ranger Arc`.

We observed that the peaks produced by the `Cell Ranger ATAC` output cover the largest number of bases across all datasets. Notably, peaks identified using `Cell Ranger ATAC` custom peak caller cover more bases than those identifed by `MACS2` [13]. On human datasets, peaks produced by `alevin-fry-atac` cover more bases compared to `Chromap`, while on mouse datasets, `Chromap` covers more bases. The peak overlap between the methods exceeds 90% across all datasets, and in most cases, it is greater than 95% (Table 2). The overlap between `Chromap` and `Cell Ranger ATAC` is slightly higher than the overlap between `alevin-fry-atac` and `Cell Ranger ATAC`— though this difference reduces substantially when using the peak range of `Cell Ranger ATAC` as the denominator.

**Table 2:** Percentage overlap between the peaks pairwise for the different methods across the datasets. Depending on which method is used as a denominator, percentage overlap can change. The values in the parenthesis denote the overlap when the second method was used as the denominator. N/A entries denote that `Cell Ranger ATAC` was not run on that dataset.

| Method / Dataset | alevin-fry-atac vs Chromap | alevin-fry-atac vs Cell Ranger ATAC | Chromap vs Cell Ranger ATAC |
| --- | --- | --- | --- |
| Human 10K PBMC | 96.59 (97.97) | 96.84 (97.02) | 99.14 (97.92) |
| Mouse 8K Cortex | 97.99 (98.31) | 97.56 (94.35) | 98.37 (94.82) |
| Human 3K Brain | 92.23 (97.80) | N/A | N/A |

#### Comparing the clusters

We performed clustering of cells for the different datasets using Signac `Signac` [36] and conducted a pairwise comparison between the clusters, both qualitatively and quantitatively. Across datasets, we observed a large overlap between clusters (Figures S14 and S15). On the `Human 10K PBMC` dataset, there was less crossover (i.e. more concordance) between the clusters obtained for `Cell Ranger ATAC` and `alevin-fry-atac` compared to other pairwise comparisons. Similarly, the UMAP plots (Figures S16 to S18) revealed very similar cluster projections.

Quantitatively, we evaluated the cluster labels (Table 3) using Adjusted Rand Index (ARI) and Normalized Mutual Information (NMI). High concordance was observed across all datasets. on the `Human 10K PBMC` dataset, ARI and NMI for the comparison between `alevin-fry-atac` and `Cell Ranger ATAC` were higher than the other pairwise comparisons. On the `Mouse 8K Cortex` dataset, ARI and NMI were highest for comparison between `alevin-fry-atac` and `Chromap`. Similarly, for `Human 3K Brain` dataset. NMI *>* 0.9 is observed between `alevin-fry-atac` and `Chromap`. Although all methods showed high concordance with close values, clusters produced using `alevin-fry-atac` peaks consistently demonstrated equal or higher concordance with clusters from `Cell Ranger ATAC` compared to those from `Chromap`.

**Table 3:**
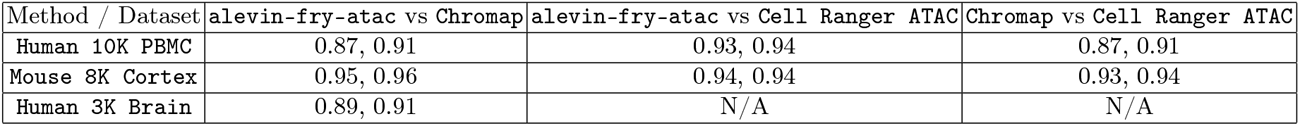
Evaluating the concordance between the clusters obtained for the different methods pairwise across the datasets. The two entries per cell separated by comma denote Adjusted Rand Index (ARI) and Normalized Mutual Information (NMI). N/A entries denote that `Cell Ranger ATAC` was not run on that dataset.

### 3.3 Memory and Running Time

Finally, we benchmarked the memory usage and runtime of `alevin-fry-atac` and `Chromap` for mapping and processing by evaluating their performance on `Human 10K PBMC` and `Mouse 8K Cortex` datasets using varying numbers of threads (Figure 3, Figure S19). Across both datasets, `Chromap` was faster than `alevin-fry-atac` when using 4 or 8 threads. However, starting at 12 threads, `alevin-fry-atac` began to outperform `Chromap`, becoming substantially faster at 32 threads, taking only 9.80 and 7.91 minutes for the `Human 10K PBMC` and `Mouse 8K Cortex` datasets respectively, where `Chromap` takes 27.45 and 21.85 minutes. Notably, `Chromap` showed poor scalability beyond 12 threads, whereas `alevin-fry-atac` demonstrated significantly improved scalability.

**Fig 3:**
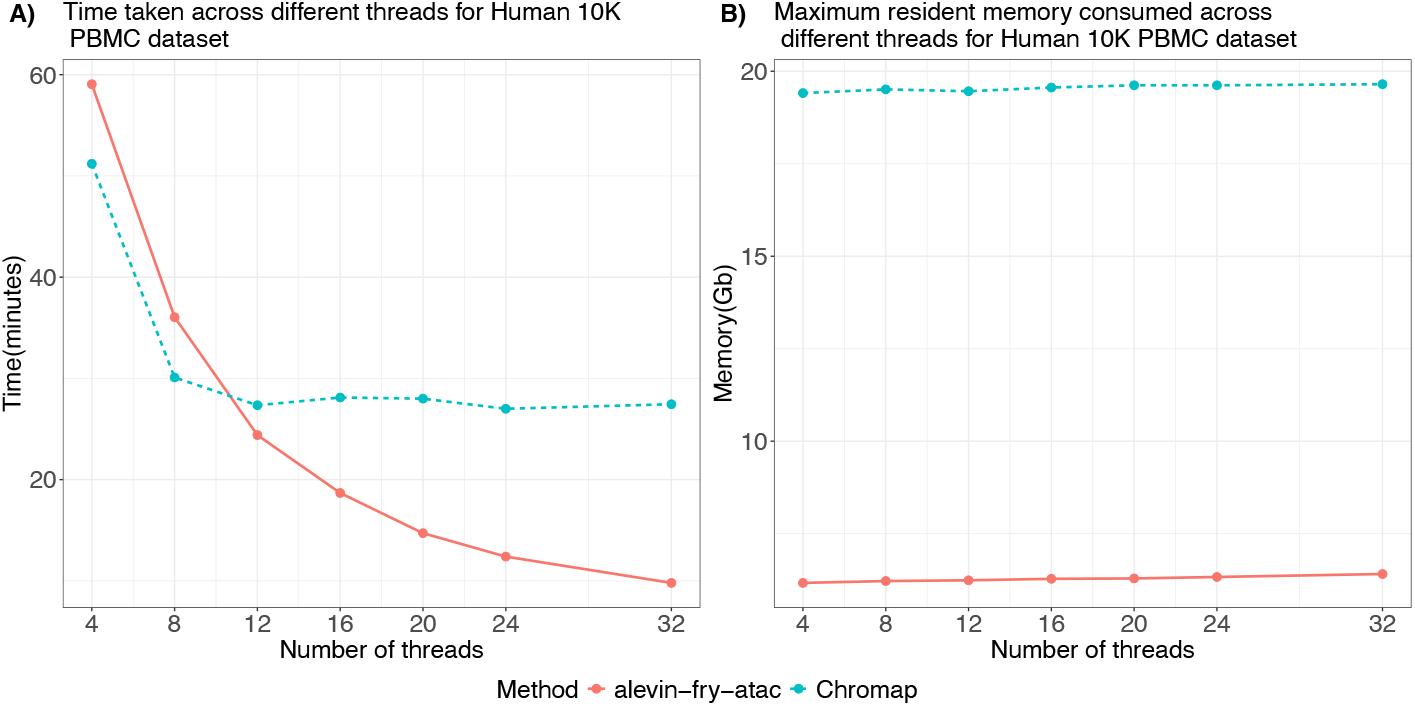
Comparing the time taken and maximum resident memory consumed by `alevin-fry-atac` and `Chromap` for the `Human 10K PBMC` dataset across the different threads.

In terms of memory usage, `alevin-fry-atac` consistently consumed substantially less memory than `Chromap`. `Alevin-fry-atac` used approximately 6.4 and 5.6 Gbs for the `Human 10K PBMC` and `Mouse 8K Cortex` datasets (Figure 3 B, Figure S19 B), compared to 19.7, 18.8 Gbs used by `Chromap`. Additionally, `Chromap`’s mapping-only mode (which only does read mapping and no other processing) and referred to as (`Chromap` (Map Only)) used 39.4, 35 Gbs for the two datassets, respectively (Figure S20, Figure S21). The benchmarking results are summarized in Table 4 for 32 threads, which also includes the results for `Cell Ranger ATAC`. However, it is important to note that the reported metrics for `Cell Ranger ATAC` also include the time spent on peak calling and other downstream analyses, as the tool does not allow for executing each step individually.

**Table 4:**
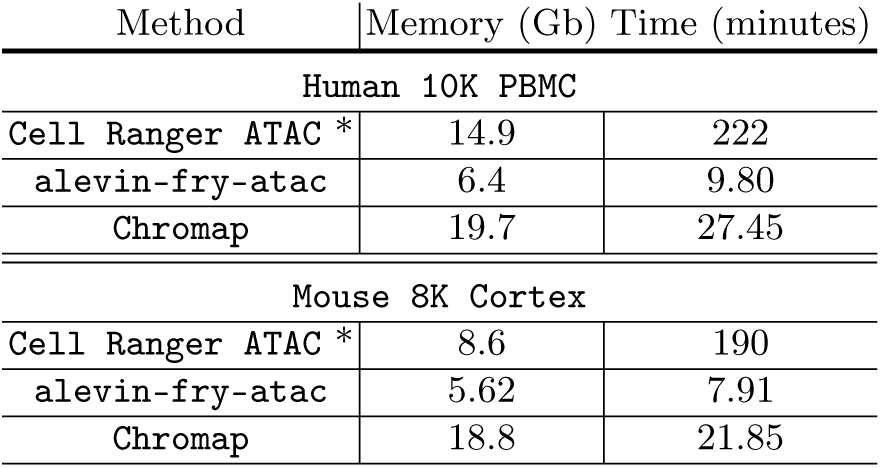
Time and maximum resident memory consumed by the different methods for mapping scATAC-seq reads. For both `Chromap` and `alevin-fry-atac` results are reported using 32 threads^1^.

| Method | Memory (Gb) | Time (minutes) |
| --- | --- | --- |
| Human 10K PBMC |  |  |
| Cell Ranger ATAC * | 14.9 | 222 |
| **alevin-fry-atac** | 6.4 | 9.80 |
| **Chromap** | 19.7 | 27.45 |
| Mouse 8K Cortex |  |  |
| Cell Ranger ATAC * | 8.6 | 190 |
| **alevin-fry-atac** | 5.62 | 7.91 |
| **Chromap** | 18.8 | 21.85 |

## 4 Discussion

We have introduced `alevin-fry-atac`, which provides a computationally efficient, memory-frugal, scalable and accurate framework for processing and mapping single-cell ATAC-seq data. It uses only ⅓ of the memory of `Chromap` and less than ½ of the memory of `Cell Ranger ATAC` when processing single-cell ATAC-seq data (and `alevin-fry-atac` utilizes only 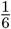of the memory of `Chromap` when the latter is not provided with a permit list during mapping). Since `Chromap` only maps the reads corresponding to the barcodes present in the permit list file, this could be one reason why we observe an increased time usage in the mapping-only mode. Internally, `alevin-fry-atac` uses the `piscem` index, which is small and versatile and enables mapping reads from other technologies such as bulk RNA-seq, single-cell RNA-seq and now single-cell ATAC-seq.

The enhanced streaming query and the introduction of cache help in significantly improving the speed and scalability of `alevin-fry-atac`. When using 12 threads, `alevin-fry-atac` outperforms `Chromap` in terms of speed and achieves even greater gains at the higher thread counts. At 32 threads, `alevin-fry-atac` processes and maps single-cell ATAC-seq data in 280% less time than `Chromap`. The introduction of virtual colors, and modifications to the pseudoalignment procedure enable accurate mapping of reads at an accuracy, achieving precision close to alignment-based methods. This level of accuracy would not have been attainable without the use of virtual colors. Despite the slightly lower mapping accuracy compared to alignment-based approaches, this does not seem to impact downstream analyses. Across experimental datasets, we observe a very high concordance between the peaks and clusters produced by `alevin-fry-atac` and those produced by the other tested methods. The `alevin-fry` and `simpleaf` ecosystem already supports the processing of single-cell RNA-seq. With the added support for single-cell ATAC-seq makes, it becomes only fully opensource software that directly supports both modalities, simplifying and facilitating multi-modal analysis. We thus believe that `alevin-fry-atac` is a viable and strong alternative to the other methods, combining efficiency, scalability, and accuracy in single-cell ATAC-seq data analysis.

While the main focus in this manuscript has been on mapping single-cell ATAC-seq data, it should be trivial to extend and apply pseudoalignment using virtual colors to other genomic-centred mapping-based tasks and modalities, such as DNase-seq, Chip-seq and Hi-C.

Finally, for our pseudoalignment strategy, we have queried all *k-*mers and utilized the “threshold-unionffi approach. However, other lightweight mapping strategies such as selective alignment [34] and pseudoalignment with structural constraints[16] can also be explored to see how they impact speed and mapping accuracy. Finally, all pipelines, including `alevin-fry-atac`, currently discard reads that map to more than one location. Such reads constitute 4 *−* 10% of the total reads in the datasets analyzed in this manuscript, and might have a non-trivial impact on downstream analysis [25]. One interesting future direction would be to develop methods to resolve the origin of such reads probabilistically using methods such as expectation-maximization [10].

## 5 Code availability

The `alevin-fry-atac` software is written in Rust and C++ and is freely available under BSD 3-Clause license, as a free and open-source tool at https://github.com/COMBINE-lab/alevin-fry-atac. `Alevin-fry-atac` has also been integrated into `simpleaf` [15], available at https://github.com/COMBINE-lab/simpleaf and via `bioconda` [14], via the `atac` sub-command.

## Footnotes

1 For `Cell Ranger ATAC` it is not possible to run and thus benchmark each step individually, thus the time and memory consumption also include other tasks such as peak calling and clustering.

