## Supplementary materials for "Alevin-fry-atac enables rapid and memory frugal mapping of single-cell ATAC-seq data using virtual colors for accurate genomic pseudoalignment"

### S1 Methods

#### S1 Mapping and Pseudoalignment

*Merging paired-end reads before mapping* — Before mapping the individual ends of a paired-end read, we try to see if we can merge the reads to create a single fragment before mapping. This idea has been suggested and adopted in several existing read-mapping tools (e.g. `STAR` [1] attempts to merge overlapping paired-end reads before mapping, and `Chromap` uses this approach for adaptor trimming [6]). We first try to merge the reads in dovetail orientation and, if that is not possible, then attempt to find an overlap merge. For a dovetail merge, the prefix of one mate should match with the suffix of the reverse complement of the other mate. For an overlap merge, the suffix of one mate should intersect with the prefix of the reverse complement of the other mate. For the merge to be successful, there should be at least an overlap of a certain number of bases between the mates (we use a default of 30). If the mates can be merged, then the merged fragment and not the individual mates are mapped to the reference.

#### S2 Analysis pipeline

The analysis pipelines used to carry out the experiments in this manuscript were created using `Snakemake` [3]. Below, we describe how the individual experiments were carried out and evaluated.

*Simulated data* — `Chromap` was run without any `--preset` arguments. For `Bowtie2`, reads were aligned by setting the maximum fragment length (`X`) to 2000. These settings were taken from `Chromap`'s experiment repository on GitHub. `Alevin-fry-atac` always maps all the records irrespective of the whitelist file. The accuracy of the mappings was evaluated using a `Python` script that was originally used for evaluation of mapping accuracy in the stobemers [4] paper.

*Experimental datasets* — `Chromap` was run with `--preset atac` argument, also taking a whitelist file as an input. `Human 3K Brain` is a multiomic dataset containing single-cell RNA-seq and single-cell ATAC-seq data. In this paper, we have only utilized single-cell ATAC-seq data for it. We ran `Cell Ranger ATAC` only on the `Human 10K PBMC` and `Mouse 8K Cortex` datasets however, since `Cell Ranger ATAC` could not process the single-cell ATAC-seq data. It required running a different software `Cell Ranger Arc`, in order to process it, which we did not run (and could not easily compare). The downstream pipelines for analyzing ATAC-seq data such as `Signac` [5] require as input a `tabix`-indexed fragment file as produced in the `Cell Ranger ATAC` pipeline. Thus, a tabix index was created for the fragment files produced by both `alevin-fry-atac` and `Chromap`. We then use `MACS2` 2 [2] as a peak caller on the fragment files produced by all the methods. The peaks and the fragment files are then passed to `Signac` for quality control and clustering.

*Benchmarking* — We benchmark both genomic read mapping and the end-to-end pipeline for processing and mapping single-cell ATAC-seq data for the different methods using `/usr/bin/time` command. For `alevin-fry-atac`, this process is divided into three steps : mapping reads to the genome, correction of barcodes from a whitelist file, and deduplication with sorting. Each steps was benchmarked individually, and their execution times were summed to calculate the total runtime for `alevin-fry-atac`. `Chromap` was benchmarked with and without using the `--preset atac` arguments. When this argument is used, then `Chromap` also performs barcode correction and sorting automatically, in addition to mapping reads to genome.

Without the argument, `Chromap` only maps reads to the genome. The methods were benchmarked across multiple thread counts, with each process run three times per thread count, and the average across the three runs was computed to obtain the final metrics (total time taken and maximum memory used). The execution time metrics for genomic read mapping process for `Chromap` (run using `--preset atac` argument), were obtained by subtracting the time taken for sorting, deduplication and writing from the total time taken for program execution. `Chromap` dumps the time for non-mapping related tasks to `stdout`. We ran `Cell Ranger ATAC` only once with 32 cores and 100 Gb of allowed RAM, since it was not possible to benchmark the individual steps. `Cell Ranger ATAC` has previously been shown to be substantially slower than `Chromap`.

*Software versions* — The following software versions were used for the analyses in this manuscript: `Cell Ranger ATAC` version 2.1.0, `Chromap` version $0.2.7 - r493$, `Bowtie2` version 2.5.4, `mason` version 2.0.9, `MACS2` version 2.2.9.1, `R` version 4.3.2, `Signac` version 1.13.0, `tabix` version 1.16, and `Snakemake` version 8.18.0. Mapping for `alevin-fry-atac` was done using `piscem` version 0.11.0 and subsequent processing was done using the `alevin-fry-atac` branch of the `alevin-fry` repository.

### S2 Results

#### S1 Mapping accuracy for `alevin-fry-atac`

We first varied the parameter $\ell_{\text{vcol}}$ (which controls the number of bases a virtual color will span, with most virtual colors spanning $\ell_{\text{vcol}} + ov\_length$ bases), for a given $k$-mer size $k$ and pseudoalignment threshold $\tau$ and observed its impact on mapping accuracy on the simulated data for the different read lengths (Figures S1 to S4). We find that accuracy decreases continuously as $\ell_{\text{vcol}}$ increases, irrespective of the values of the other parameters. Importantly, the sharpest decrease is observed when the chromosomes themselves are used as the colors for pseudoalignment (removing the effect of virtual colors on the procedure entirely). The accuracy drops to $84-86\%$ from being $92-96\%$ (depending on the read length) when using virtual colors. In general, we observe a drop in accuracy of $\sim 10\%$ or more when disabling virtual colors. Longer reads generally yield higher accuracy across most settings, except in the specific parameter configuration where $\tau = 1$ and virtual colors are disabled, where reads with length 100 have the highest accuracy.

We next varied the threshold $\tau$ while keeping $\ell_{\text{vcol}}$ and $k$ fixed (Figures S5 to S8). The accuracy decreases as the threshold decreases from 0.6 to 1. The decline seems to be sharpest from 0.8 to 1, with the magnitude of decrease in accuracy in general being much higher for larger read lengths. For $\tau = 1$, at all the $\ell_{\text{vcol}}$ values, the accuracy of reads with length 150 is smaller than reads with length 100. However, when the chromosomes are used as colors, the accuracy for reads with length 150 increases when $\tau$ increases from $0.6 - 0.8$ and falls again at 1.

Finally, we varied $k$, while keeping $\ell_{\text{vcol}}, \tau$ fixed (Figures S9 to S12). We observe that for reads of length 50, accuracy increases from $k = 23$ to $k = 25$ and starts decreasing across most virtual color extends and thresholds. However, when chromosomes are used as colors, accuracy increases with an increase in $k$. For the reads with larger lengths, a larger $k$ is preferred as the accuracy increases uniformly across parameters.

### S3 Tables

Table S1: URLs for the datasets used in the manuscript for analysis.

| Dataset | URL |
|---|---|
| `Human 10K PBMC` | 10K |
| `Mouse 8K Cortex` | 8k |
| `Human 3K Brain` | 3K |

Table S2: The total number of bases (MB) covered by the different methods across the different datasets. N/A entries denotes that `Cell Ranger ATAC` was not run on that dataset. `Cell Ranger ATAC` (`MACS2`) and `Cell Ranger ATAC` (Custom) denote that peaks were generated on the `Cell Ranger ATAC` fragments using `MACS2` and the inbuilt peak caller of `Cell Ranger ATAC` respectively.

| Method/ Dataset | `alevin-fry-atac` | `Chromap` | `Cell Ranger ATAC` (`MACS2`) | `Cell Ranger ATAC` (Custom) |
|---|---|---|---|---|
| `Human 10K PBMC` | 98.33 | 97.42 | 98.60 | 140.18 |
| `Mouse 8K Cortex` | 107.86 | 108.32 | 112.18 | 150.80 |
| `Human 3K Brain` | 115.14 | 111.80 | N/A | N/A |

### S4 Figures

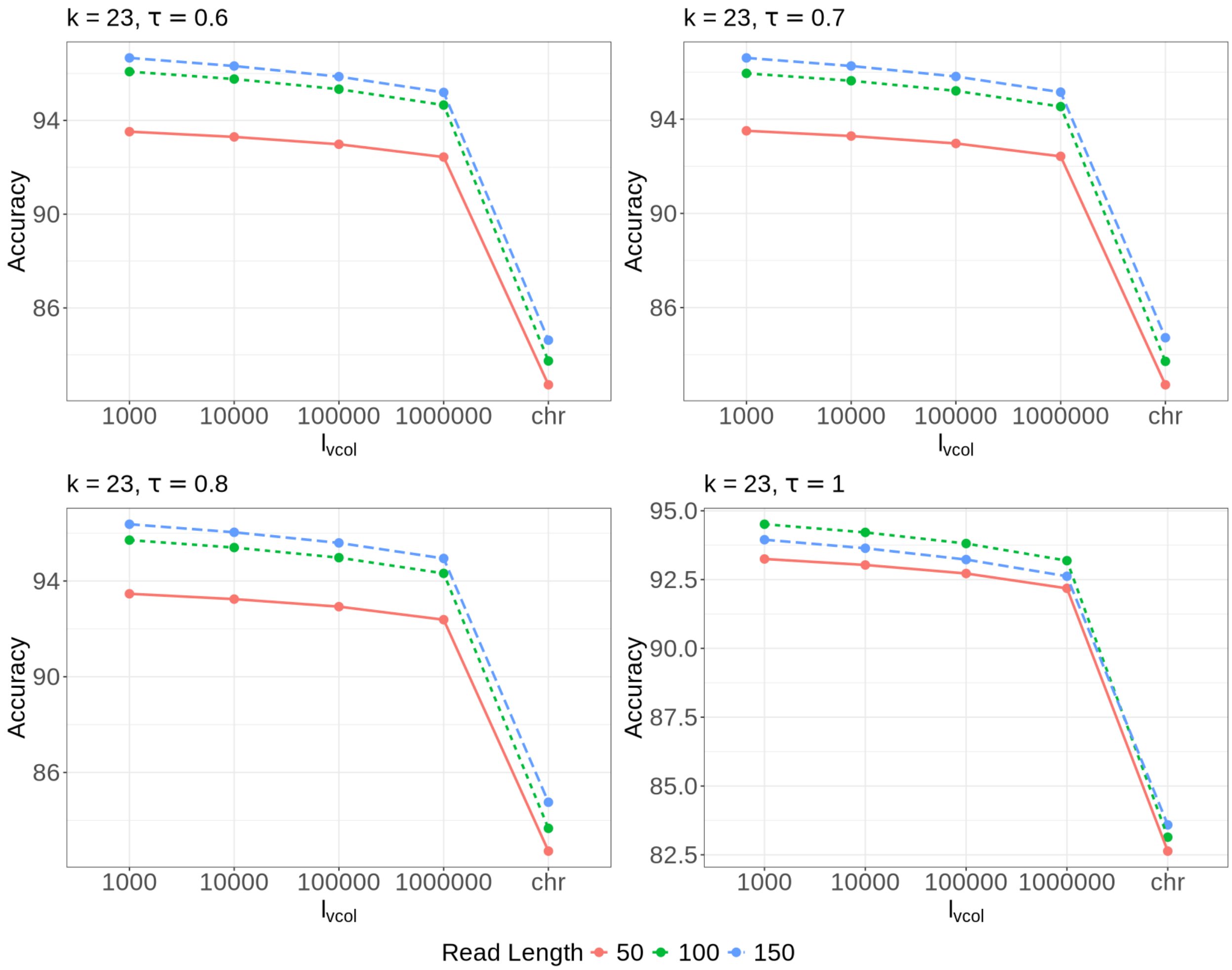


Fig. S1: Evaluating accuracy across different read lengths for `alevin-fry-atac` by varying $\ell_{\text{vcol}}$ at $k = 23$, with panels representing different $\tau$ values. The label *chr* on the x-axis implies mapping is done using chromosome sequence as colors (removing the impact of virtual colors on mapping procedure).

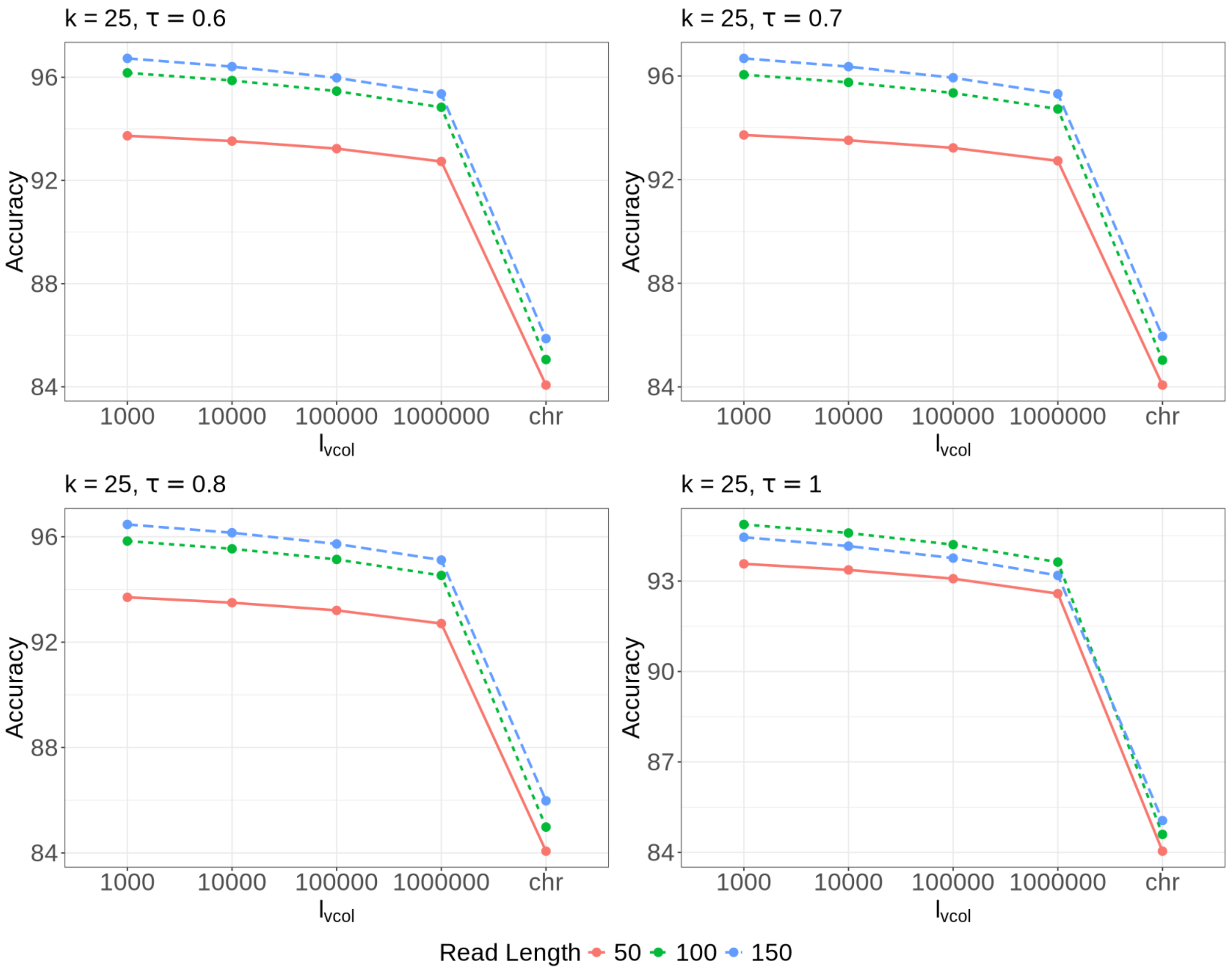


Fig. S2: Evaluating accuracy across different read lengths for `alevin-fry-atac` by varying $\ell_{\mathrm{vcol}}$ at $k = 25$, with panels representing different $\tau$ values. The label *chr* on the x-axis implies mapping is done using chromosome sequence as colors (removing the impact of virtual colors on mapping procedure).

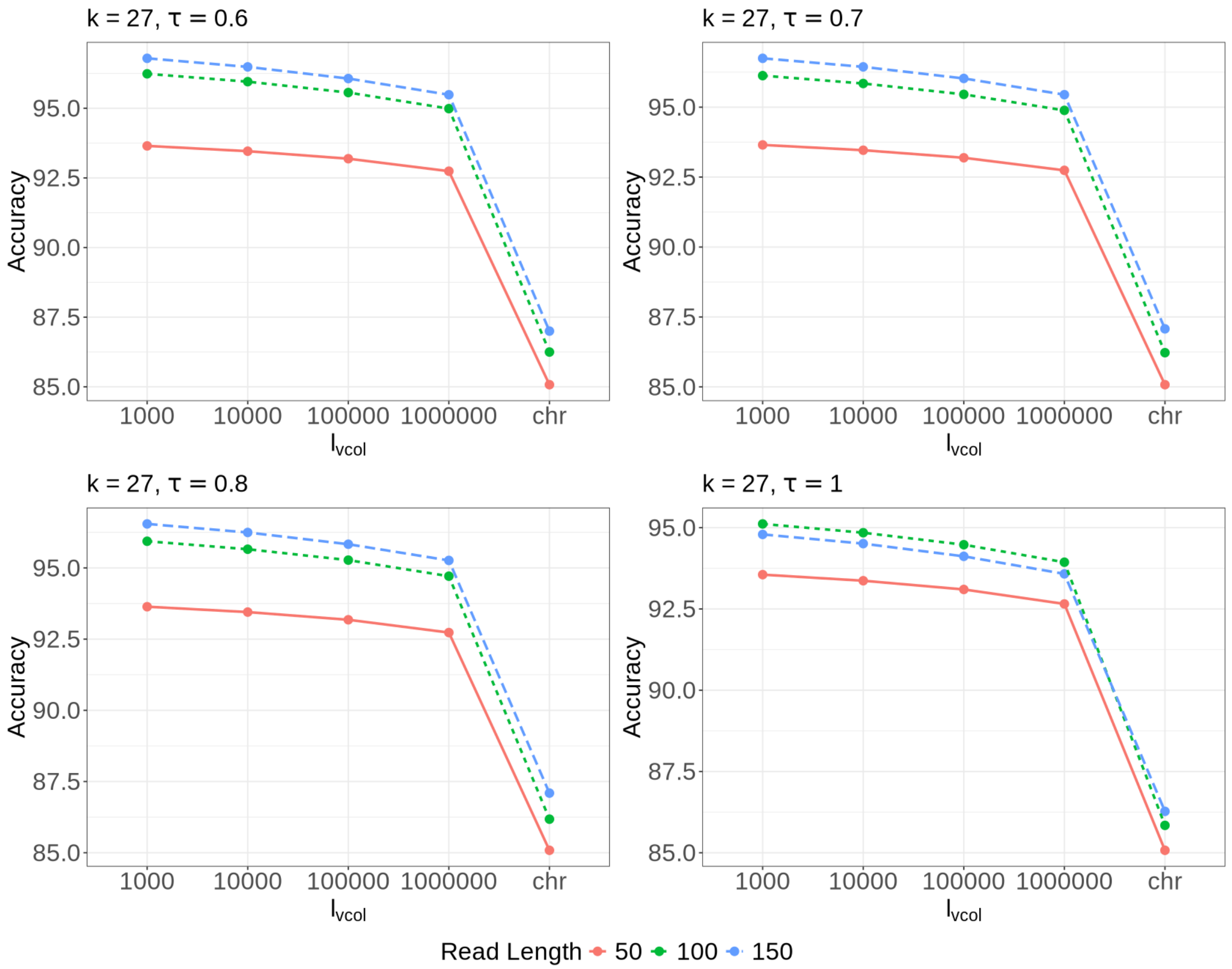


Fig. S3: Evaluating accuracy across different read lengths for `alevin-fry-atac` by varying $\ell_{\text{vcol}}$ for $k = 27$ with panels representing different $\tau$ values. The label *chr* on the x-axis implies mapping is done using chromosome sequence as colors (removing the impact of virtual colors on mapping procedure).

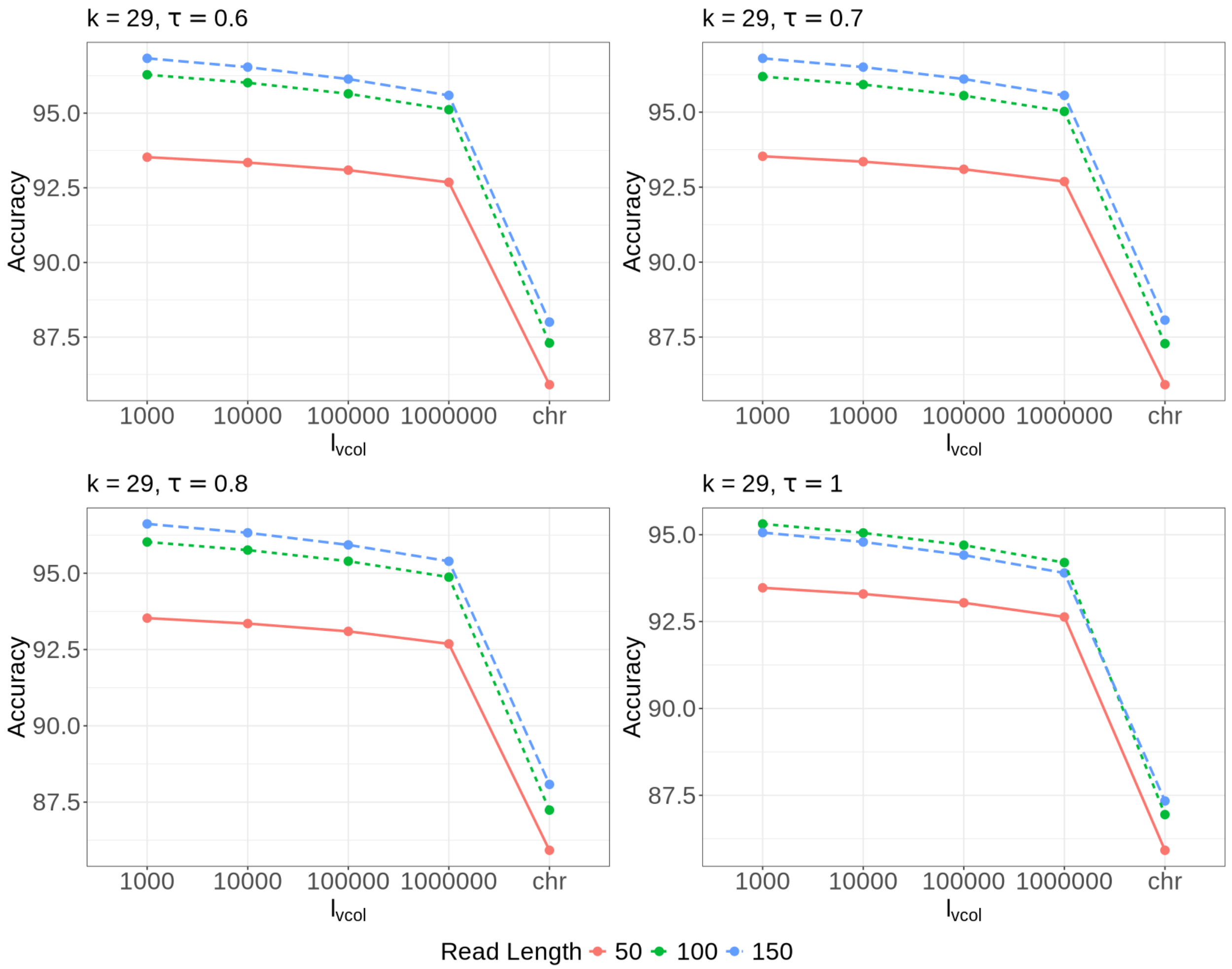


Fig. S4: Evaluating accuracy across different read lengths for `alevin-fry-atac` by varying $\ell_{\text{vcol}}$ for $k = 29$ with panels representing different $\tau$ values. The label *chr* on the x-axis implies mapping is done using chromosome sequence as colors (removing the impact of virtual colors on mapping procedure).

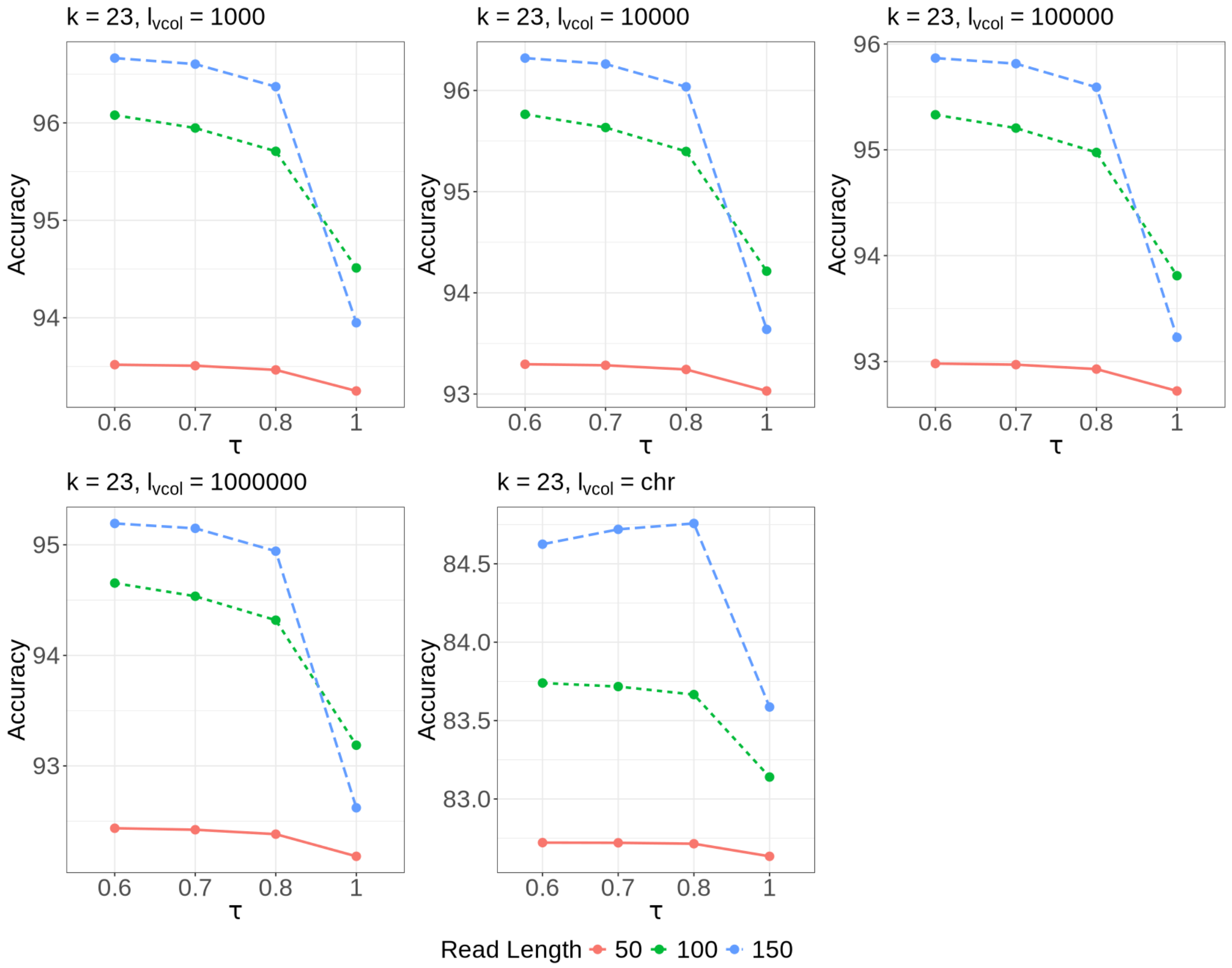


Fig. S5: Evaluating accuracy across different read lengths for `alevin-fry-atac` by varying the pseudoalignment thresholds for $k = 23$ with panels representing different $\ell_{\text{vcol}}$ values. The panel with $\ell_{\text{vcol}}$ 1$chr$ implies mapping is done using chromosome sequence as colors (removing the impact of virtual colors on mapping procedure).

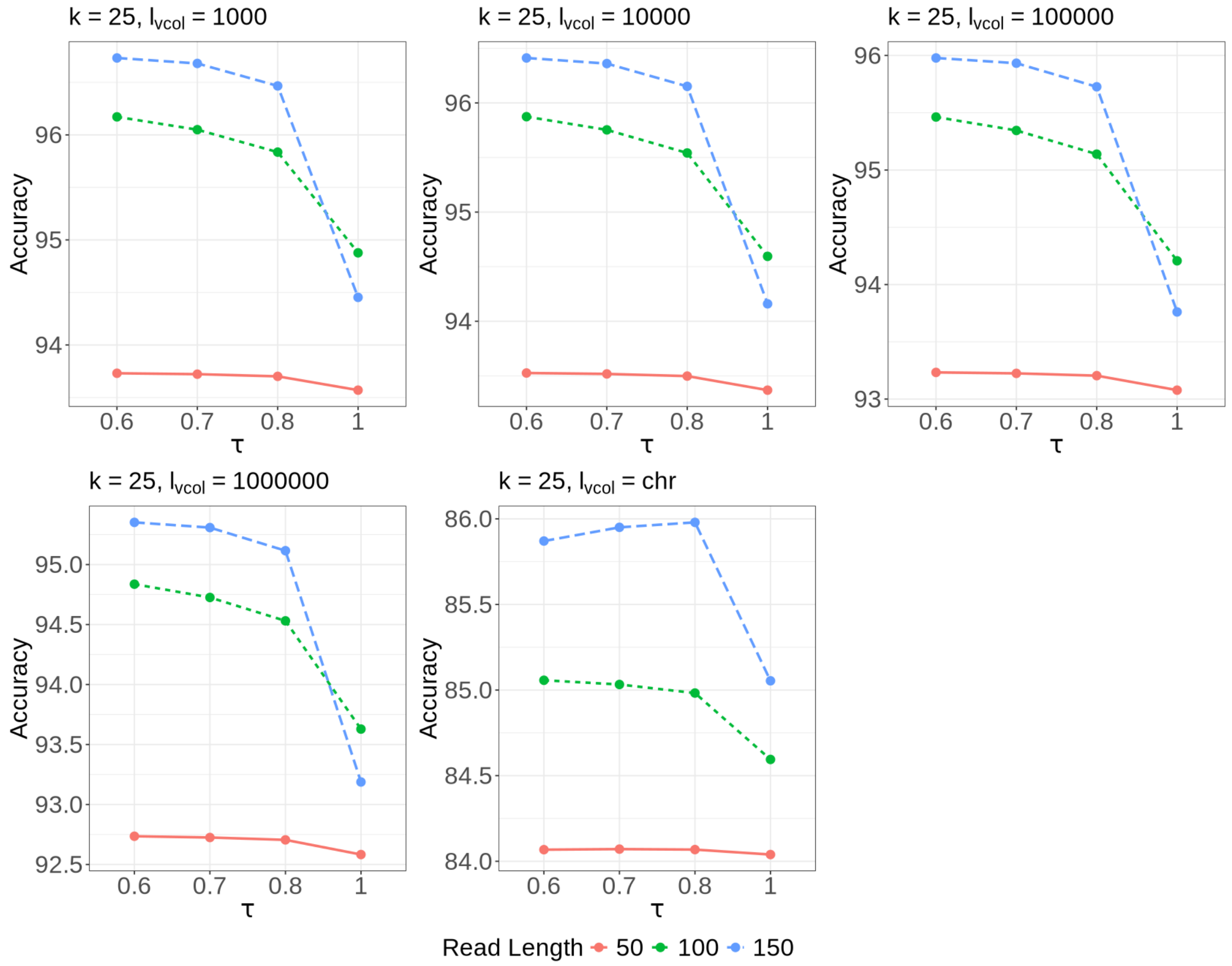


Fig. S6: Evaluating accuracy across different read lengths for `alevin-fry-atac` by varying the pseudoalignment thresholds for $k = 25$ with panels representing different $\ell_{\mathrm{vcol}}$ values. The panel with $\ell_{\mathrm{vcol}}$ *chr* implies mapping is done using chromosome sequence as colors (removing the impact of virtual colors on mapping procedure).

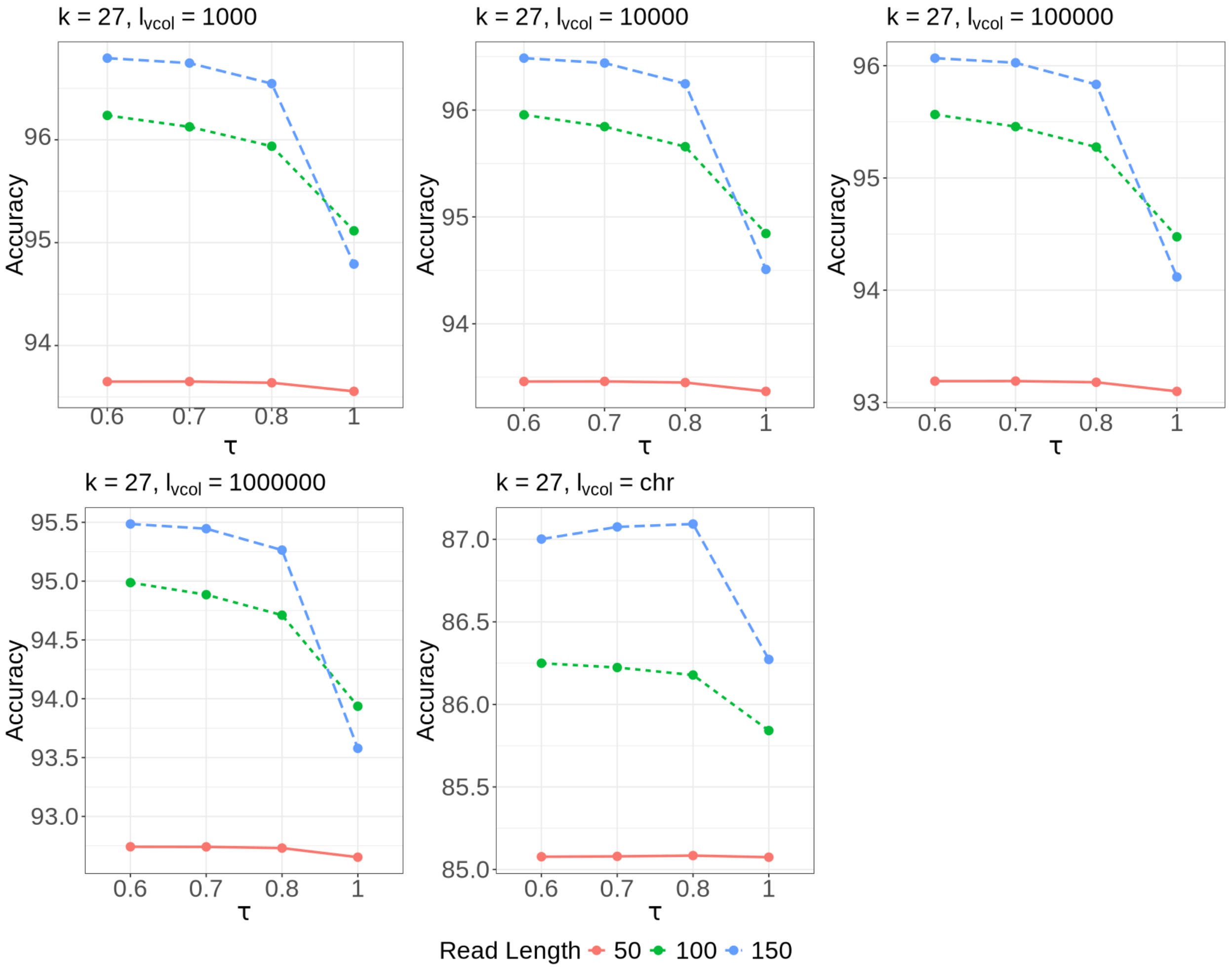


Fig. S7: Evaluating accuracy across different read lengths for `alevin-fry-atac` by varying the pseudoalignment thresholds for $k = 27$ with panels representing different $\ell_{\text{vcol}}$ values. The panel with $\ell_{\text{vcol}}$ *chr* implies mapping is done using chromosome sequence as colors (removing the impact of virtual colors on mapping procedure).

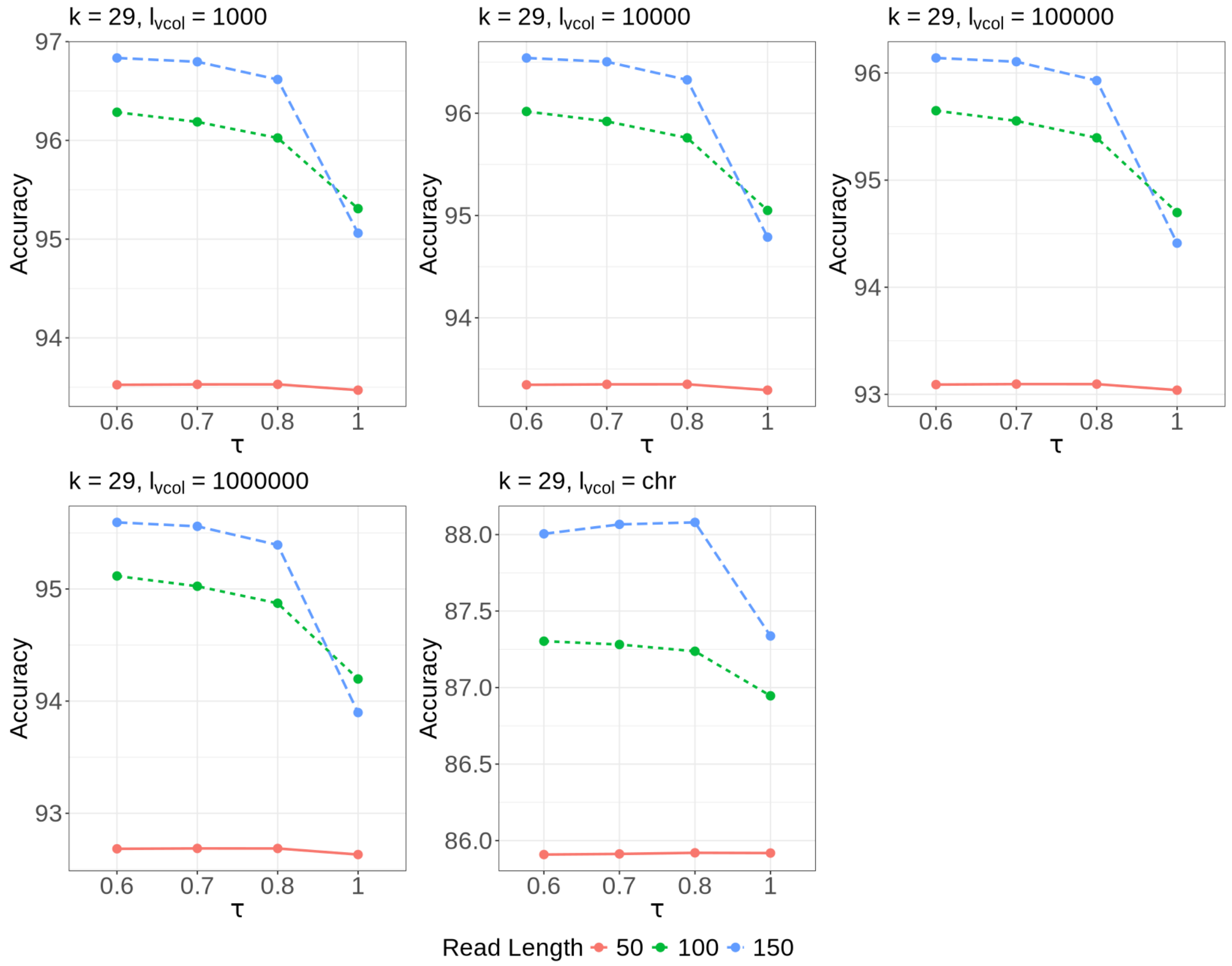


Fig. S8: Evaluating accuracy across different read lengths for `alevin-fry-atac` by varying the pseudoalignment thresholds for $k = 29$ with panels representing different $\ell_{\text{vcol}}$ values. The panel with bin length *chr* implies mapping is done using chromosome sequence as colors (removing the impact of virtual colors on mapping procedure).

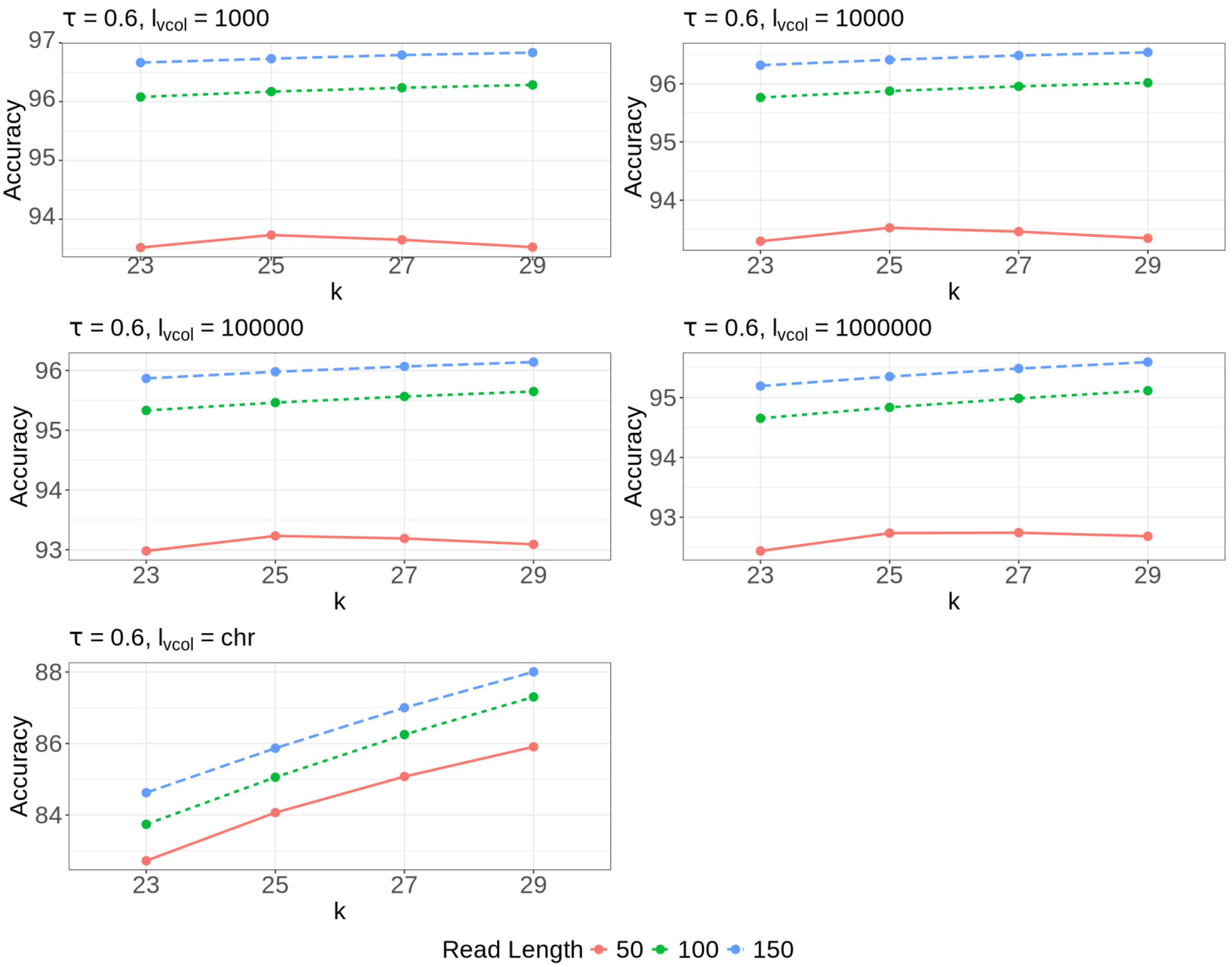


Fig. S9: Evaluating accuracy across different read lengths for `alevin-fry-atac` by varying the $k$-mer size for $\tau = 0.6$ with panels representing different $\ell_{\mathrm{vcol}}$ values. The panel with $\ell_{\mathrm{vcol}}$ *chr* implies mapping is done using chromosome sequence as colors (removing the impact of virtual colors on mapping procedure).

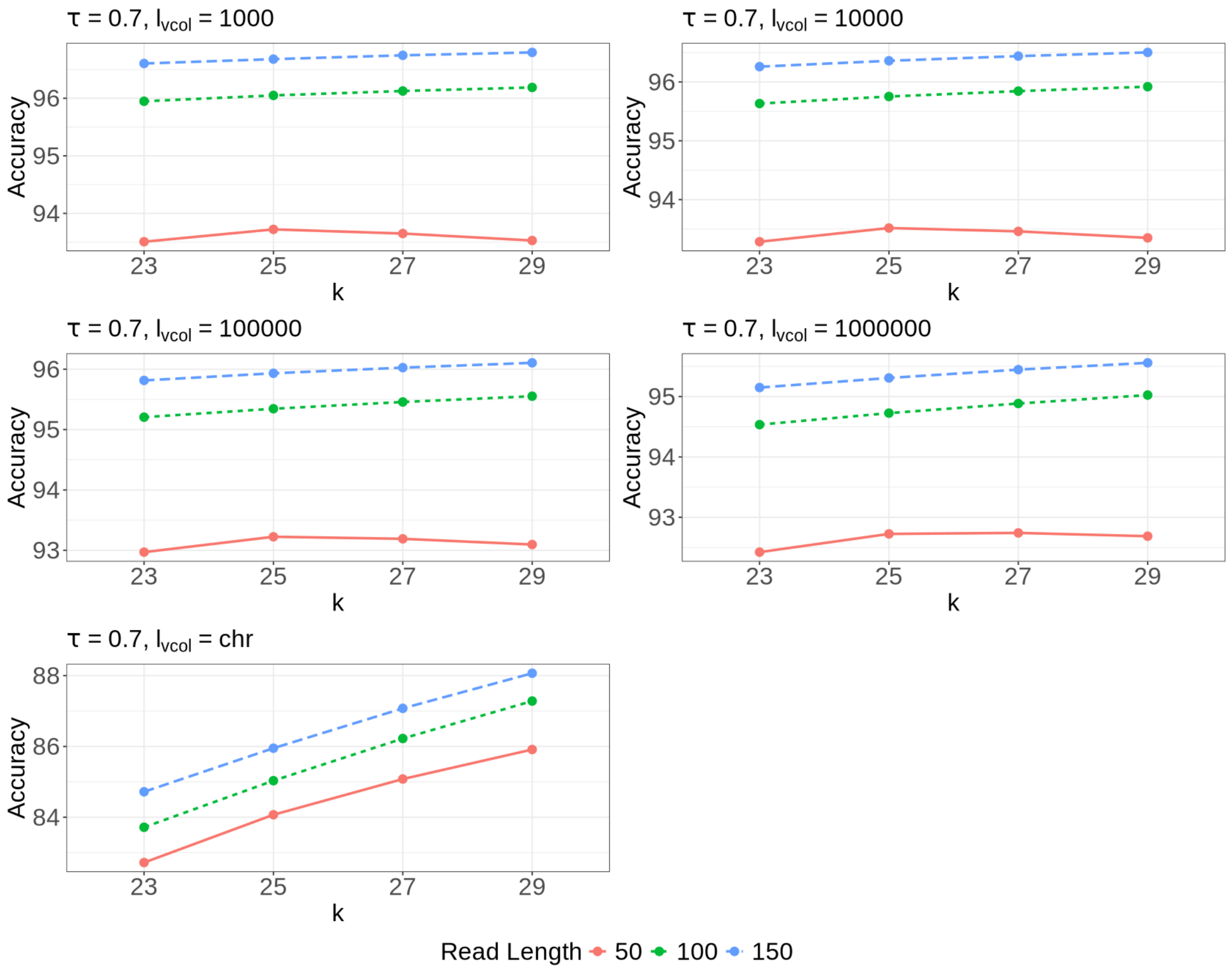


Fig. S10: Evaluating accuracy across different read lengths for `alevin-fry-atac` by varying the $k$-mer size for $\tau = 0.7$ with panels representing different $\ell_{\mathrm{vcol}}$ values. The panel with $\ell_{\mathrm{vcol}}$ *chr* implies mapping is done using chromosome sequence as colors (removing the impact of virtual colors on mapping procedure).

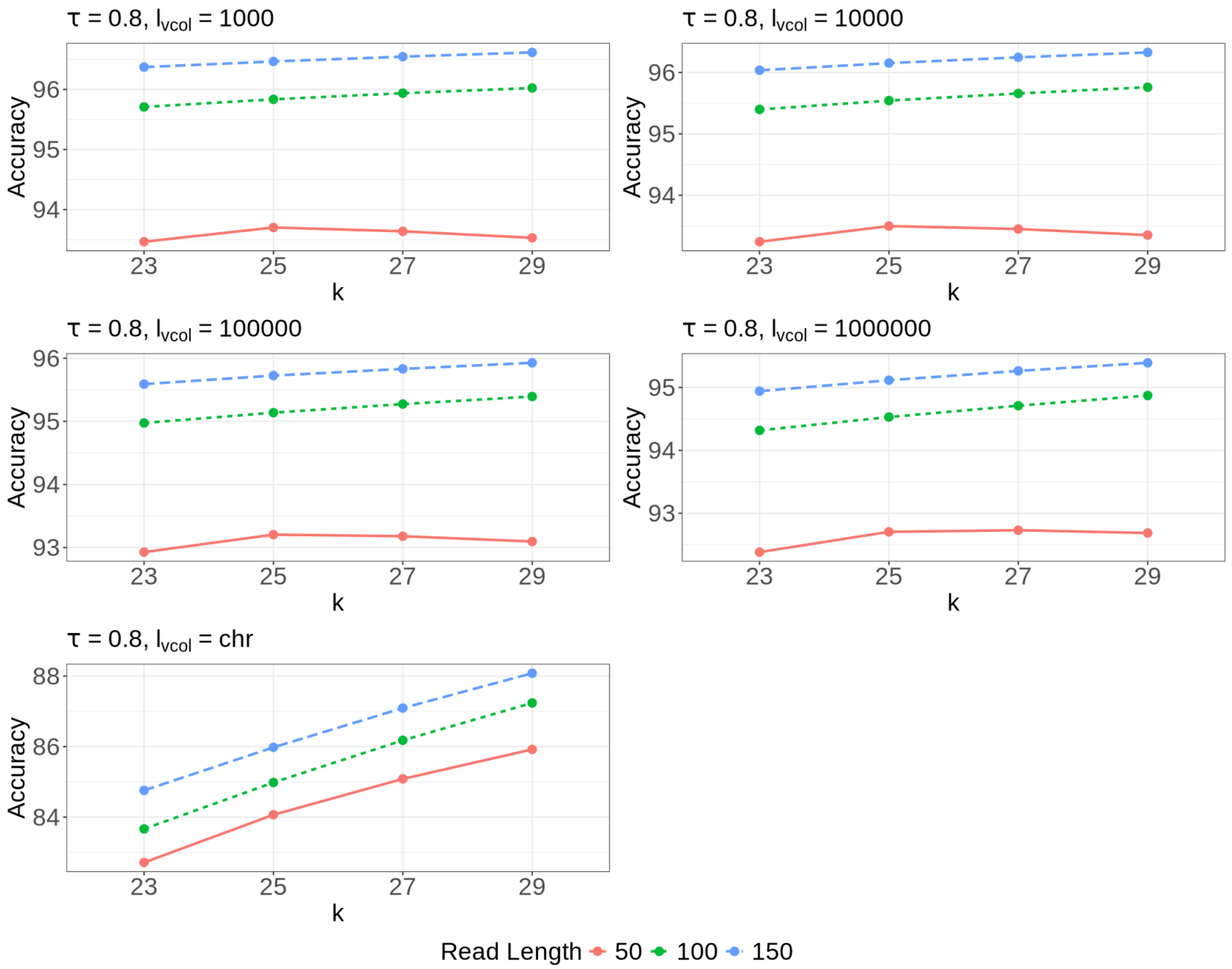


Fig. S11: Evaluating accuracy across different read lengths for `alevin-fry-atac` by varying the $k$-mer size for $\tau = 0.8$ with panels representing different $\ell_{\text{vcol}}$ values. The panel with $\ell_{\text{vcol}}$ *chr* implies mapping is done using chromosome sequence as colors (removing the impact of virtual colors on mapping procedure).

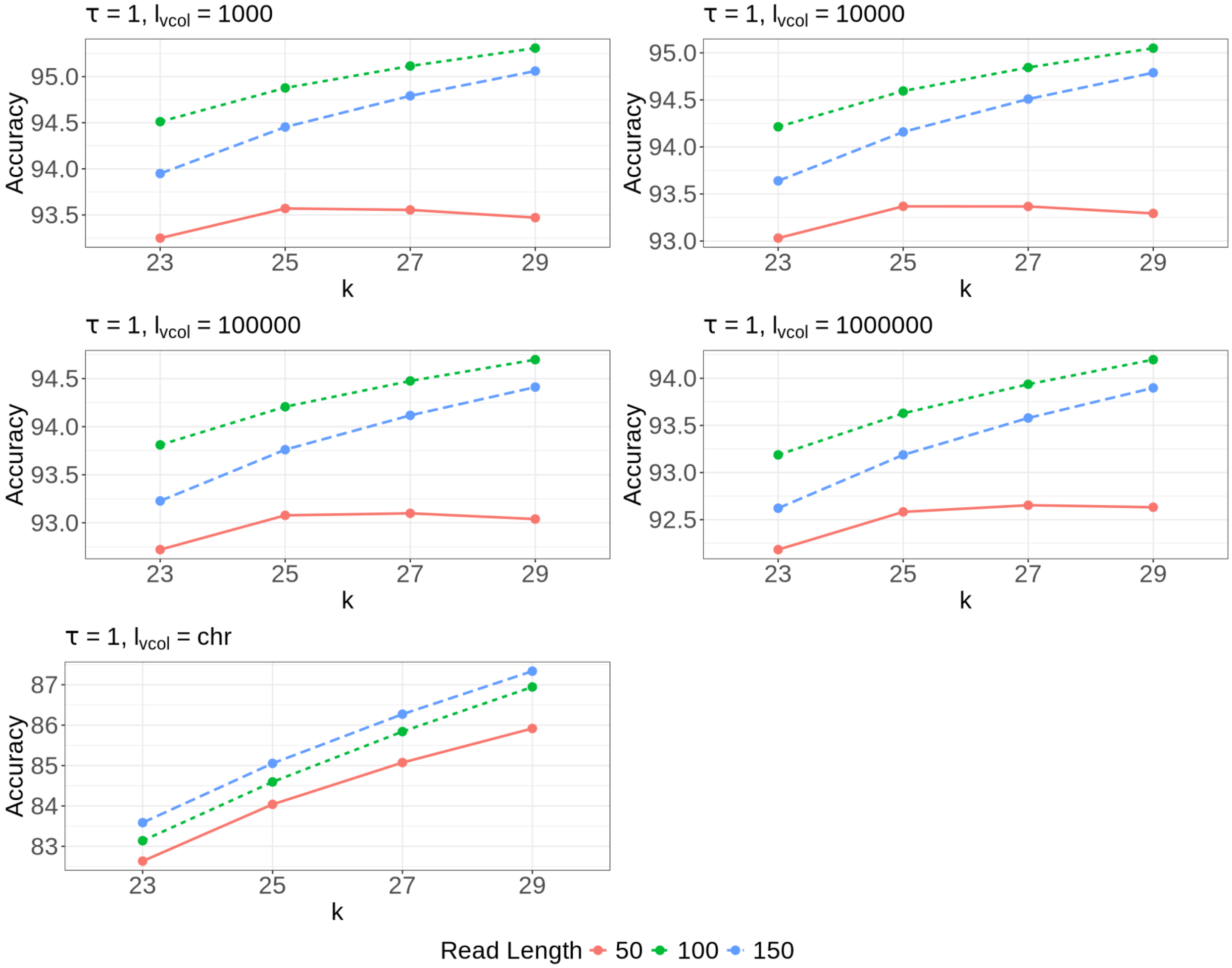


Fig. S12: Evaluating accuracy across different read lengths for `alevin-fry-atac` by varying the $k$-mer size for $\tau = 1$ with panels representing different $\ell_{\text{vcol}}$ values. The panel with $\ell_{\text{vcol}}$ *chr* implies mapping is done using chromosome sequence as colors (removing the impact of virtual colors on mapping procedure).

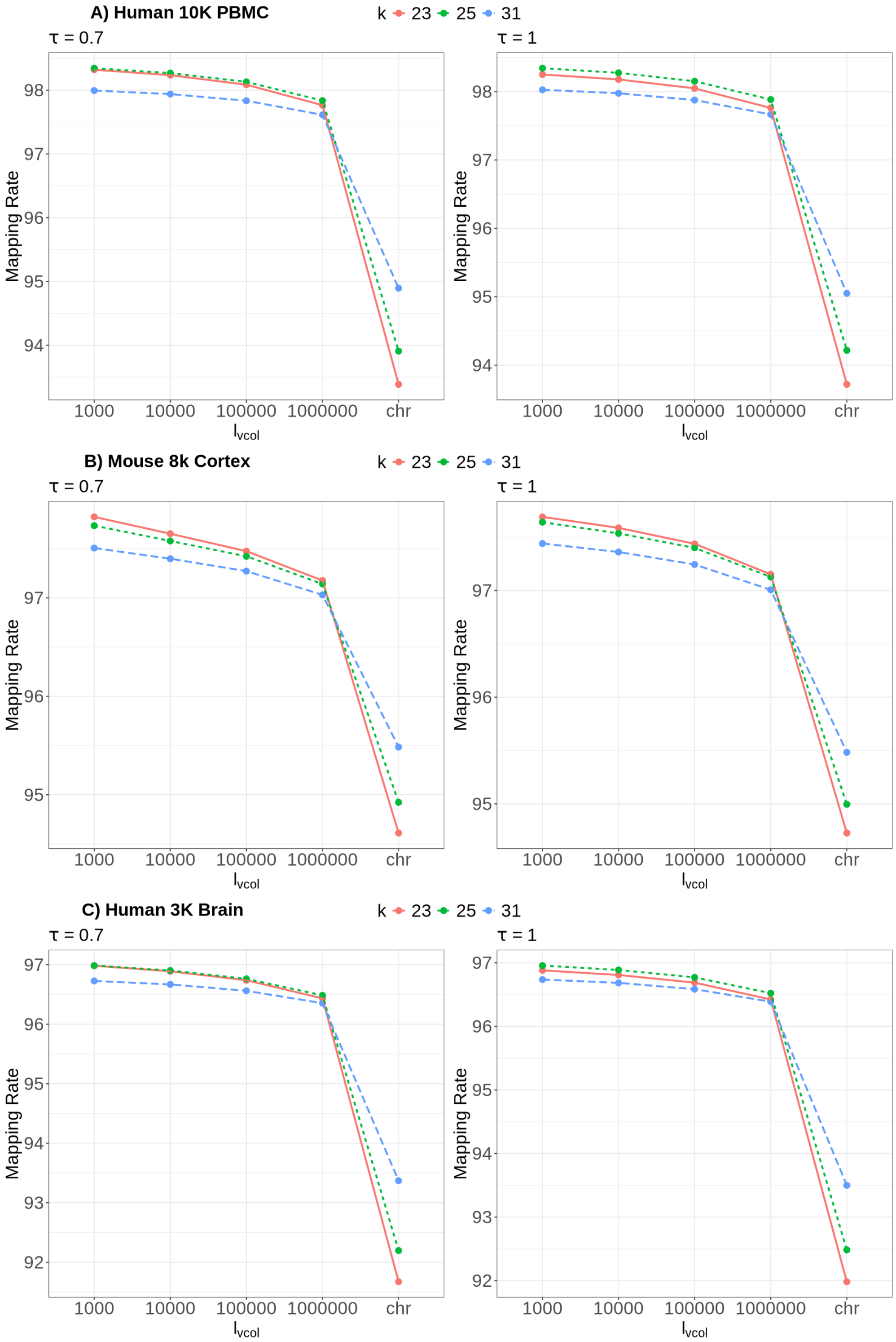


Fig. S13: Evaluating the mapping rate across different $k$-mer size for `alevin-fry-atac` by varying $\ell_{\mathrm{vcol}}$ for A) $\tau = 0.7$, B) $\tau = 1$ on the `Human 10K PBMC`, `Mouse 8K Cortex` and `Human 3K Brain` datasets respectively. The x-axis labels *chr* implies mapping is done using chromosome sequence as colors (removing the impact of virtual colors on mapping procedure).

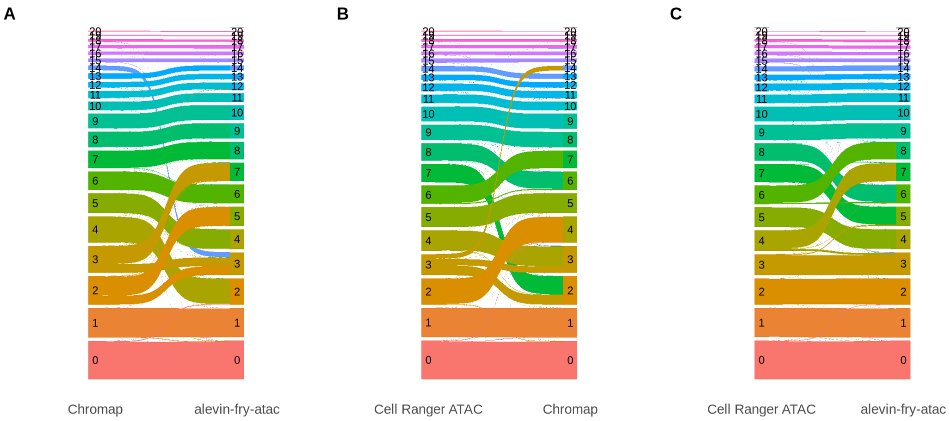


Fig. S14: Pairwise Sankey plot on the clusters obtained across the different methods for `Human 10K PBMC` dataset.

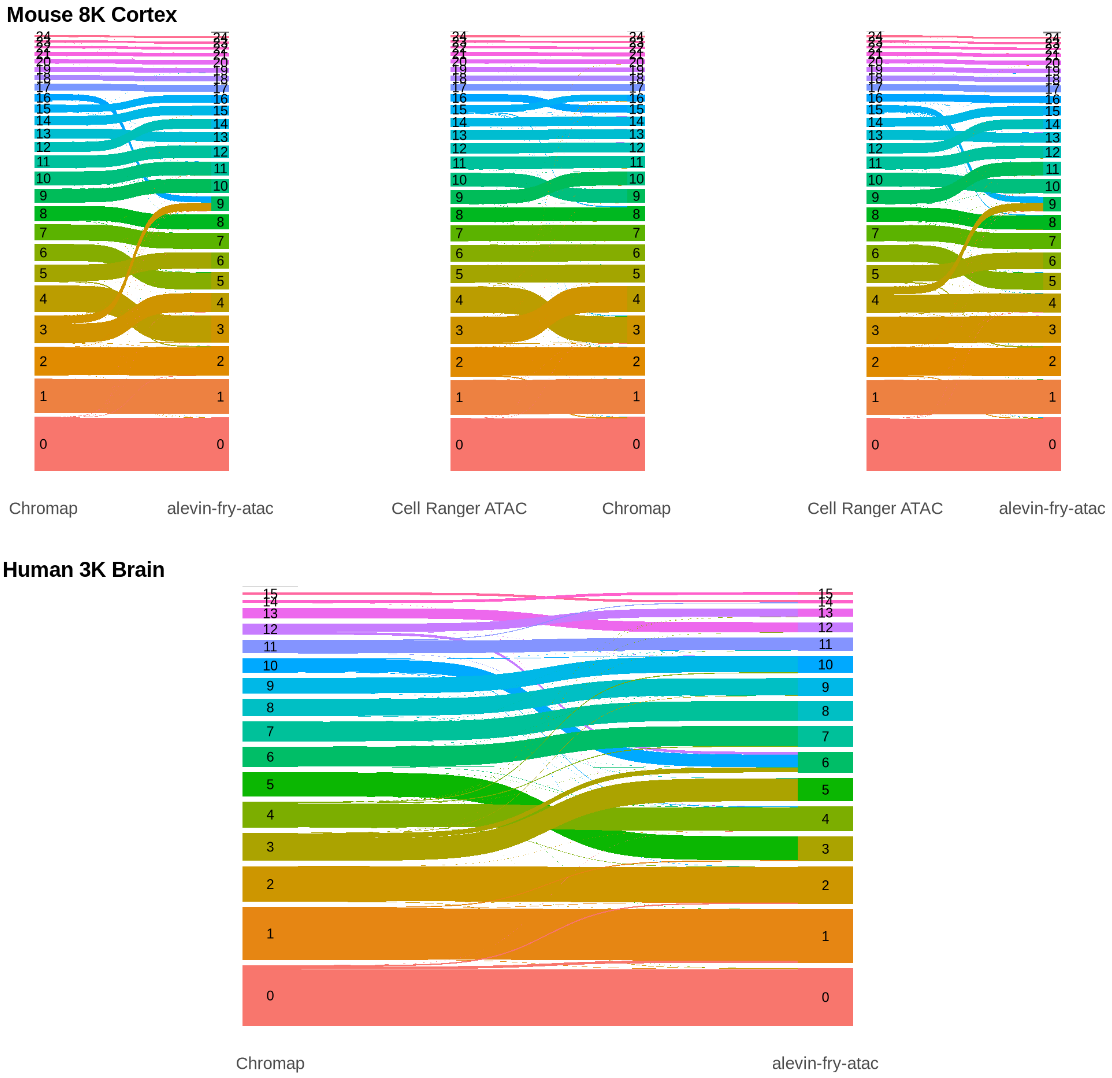


Fig. S15: Pairwise Sankey plot on the clusters obtained across the different methods for `Mouse 8K Cortex` and `Human 3K Brain` dataset.

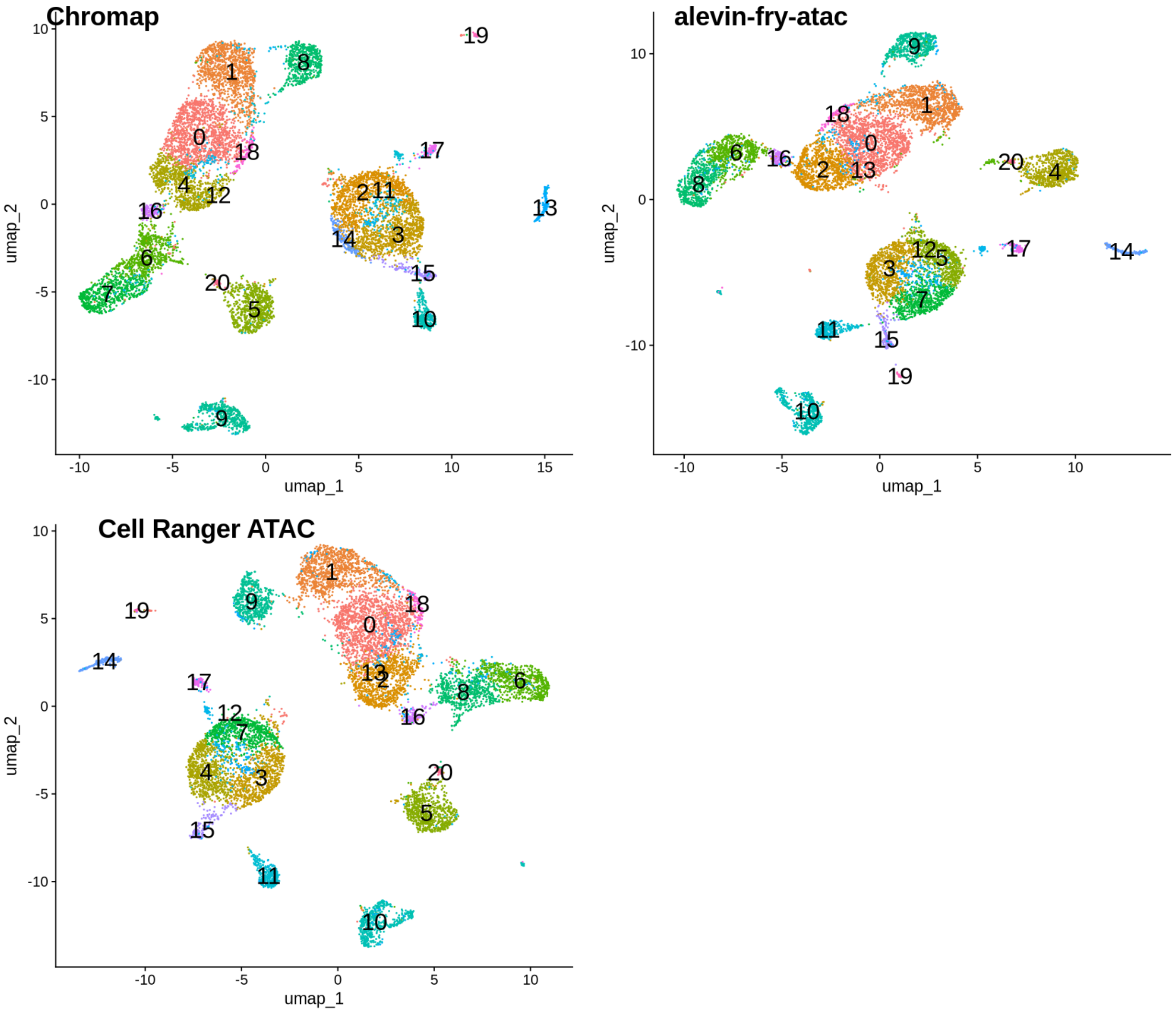


Fig. S16: UMAP projections with the cluster labels obtained for the different methods on the `Human 10K PBMC` dataset.

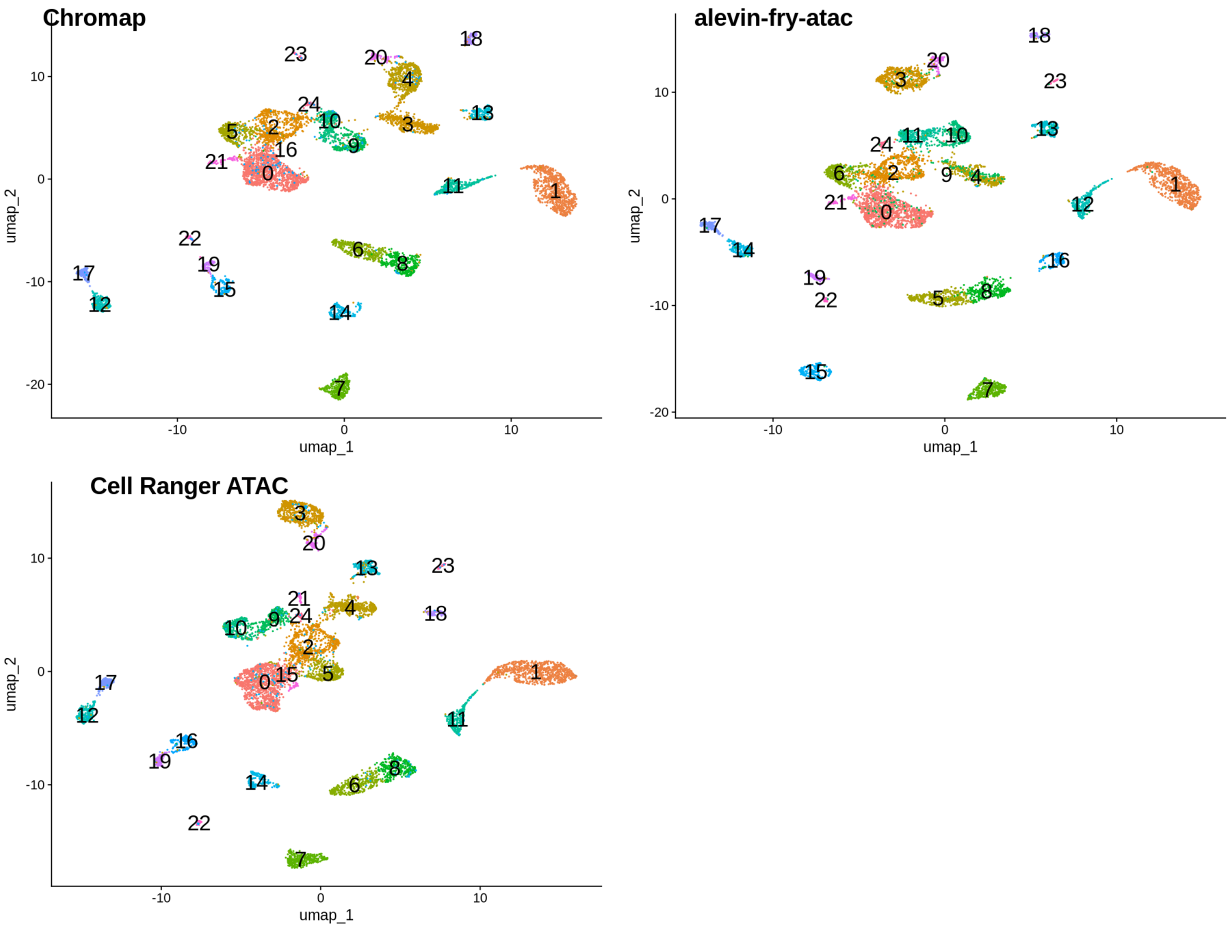


Fig. S17: UMAP projections with the cluster labels obtained for the different methods on the `Mouse 8K Cortex` dataset.

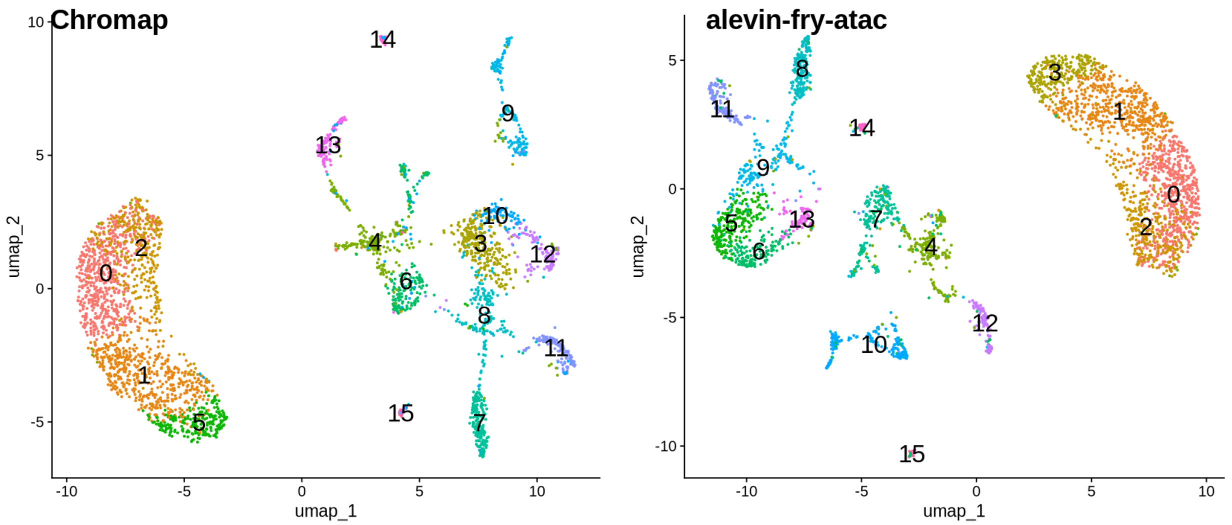


Fig. S18: UMAP projections with the cluster labels obtained for the different methods on the `Human 3K Brain` dataset.

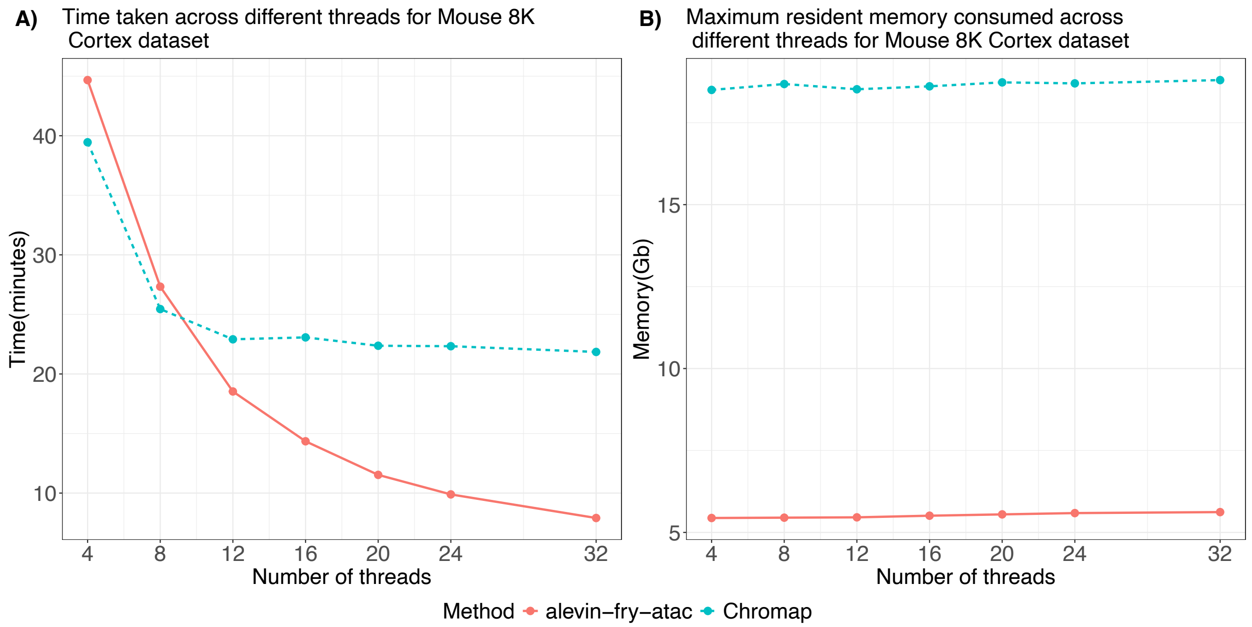


Fig. S19: Comparing the time taken and maximum resident memory consumed by `alevin-fry-atac` and `Chromap` for the `Mouse 8K Cortex` dataset across the different threads.

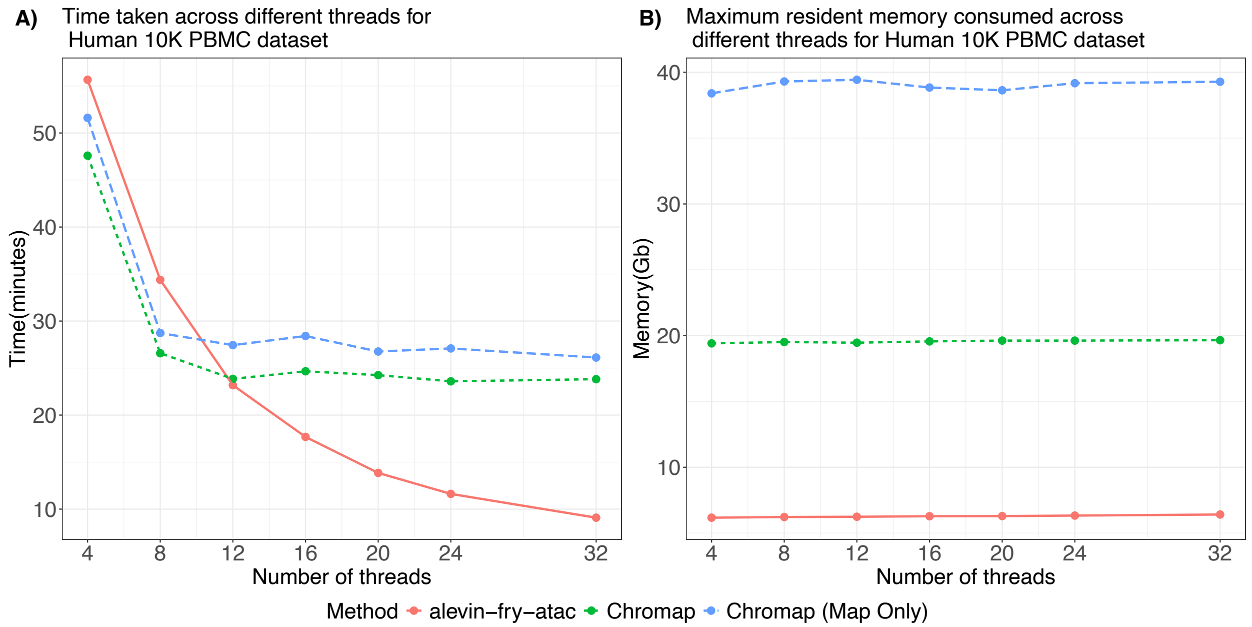


Fig. S20: Comparing the time taken and maximum resident memory consumed by `alevin-fry-atac` and `Chromap` for the `Human 10K PBMC` dataset across the different threads to map the reads. `Chromap` (Map Only) refers to when `Chromap` is run wihout the `-preset` arguments.

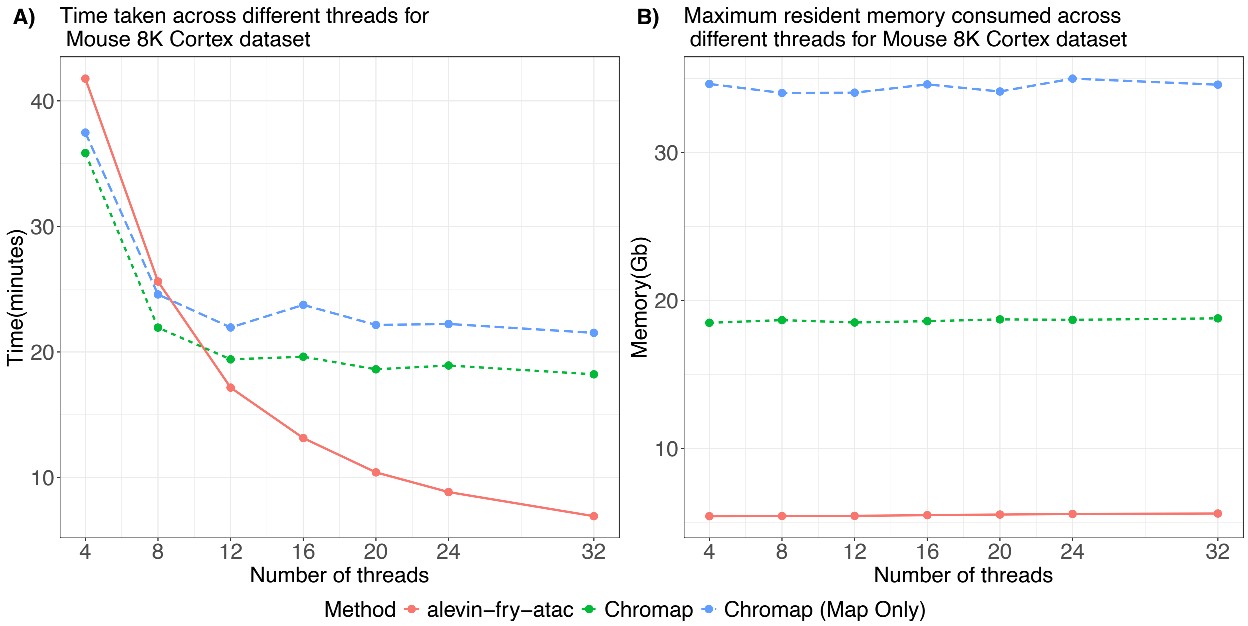


Fig. S21: Comparing the time taken and maximum resident memory consumed by `alevin-fry-atac` and `Chromap` for the `Mouse 8K Cortex` dataset across the different threads to map the reads. `Chromap` (Map Only) refers to when `Chromap` is run wihout the `-preset` arguments.
